# APP metabolism is regulated by p97/VCP through an autophagy and endolysosome-dependent mechanism

**DOI:** 10.64898/2026.02.03.703492

**Authors:** Adriana Figueroa-Garcia, Lisa Brisoire, Catherine Baud, Raphaelle Caillierez, Sabiha Eddarkaoui, Caroline Evrard, Jean-Michel Saliou, Séverine Bégard, Mathilde Coevoet, Hanane Abouelfarah, Alexandre Trichies, Luc Buée, Patricia Melnyk, Valérie Vingtdeux, Raphaelle Pardossi-Piquard, Frédéric Checler, Nicolas Sergeant

**Affiliations:** Univ. Lille, Inserm, CHU Lille, U1172—LilNCog—Lille Neuroscience & Cognition, Lille Neuro & Chemistry team, F-59000 Lille, France; Univ. Lille, Inserm, CHU Lille, U1172—LilNCog—Lille Neuroscience & Cognition, Lille Neurodegeneration team, F-59000 Lille, France; Univ. Lille, CNRS, Inserm, CHU Lille, Pasteur Institute of Lille, UMS 2014 – US41, Proteomics and modified peptides platform (P3M), Joint Service Unit PLBS (Platforms Lille in Biology & Health), Lille, France; Univ. Lille, CNRS, Inserm, CHU de Lille, Institut Pasteur de Lille, US 41 - UAR 2014 - PLBS - Plateformes Lilloises de Biologie & Santé, F-59000 Lille, France; Institut of Molecular and Cellular Pharmacology, LabEx DistAlz, Université Côte d’Azur, INSERM, CNRS, Sophia-Antipolis, 06560 Valbonne, France

**Keywords:** Alzheimer’s disease, amyloid precursor protein, valosin-containing protein, protein homeostasis, autophagy, proteasome

## Abstract

Deregulation of amyloid precursor protein (APP) metabolism leads to the production of pathological proteoforms of Aβ peptides, which ultimately form amyloid deposits, a primary pathological feature of Alzheimer’s disease (AD). The accumulation of misfolded proteins and alterations in protein degradation systems, such as the ubiquitin-proteasome system, endoplasmic reticulum-associated degradation, and autophagy-lysosomal pathways, also contribute to AD development. The Valosin-Containing Protein AAA-ATPase (p97/VCP) is a crucial regulator of proteostasis, facilitating the clearance of misfolded proteins by the proteasome and other protein degradation systems. Here, we investigated whether VCP influences APP processing. Reducing VCP expression or inhibiting its ATPase activity led to the accumulation of mature, full-length APP in cells, thereby decreasing APP trafficking to the cell surface. Downstream of APP-CTFs’ secretase processing, p97/VCP modulates APP-CTFs cleavage and degradation via autophagy and endolysosomal-dependent mechanisms. Our findings demonstrate that VCP is involved in APP metabolism at two levels: controlling APP trafficking within the cell secretory pathway and regulating autophagy-dependent degradation of APP-CTFs, suggesting a potential role for VCP in the APP deregulation observed in AD.

## 1. Introduction

Alzheimer’s disease (AD) is the most common neurodegenerative disorder and is driven by two major pathological processes: amyloid parenchymal deposition and neurofibrillary degeneration. These lesions consist of amyloid-β (Aβ) peptides and fibrillary aggregates of abnormally modified Tau proteins, respectively. Their interplay, along with neuroinflammation and genetic risk factors, defines the overall pathophysiology of the disease (Scheltens et al., 2021). A growing body of evidence suggests that amyloid deposition and neurofibrillary degeneration may be driven by a prion-like mechanism (Mudher et al., 2017; Vingtdeux et al., 2012), in which pathologically misfolded proteins propagate through the induction of trans-conformational changes in their native counterparts, ultimately leading to protein aggregation (Duykaerts et al., 2019).

Cellular proteostasis systems progressively decline with aging and in neurodegenerative diseases (Hipp et al., 2019). These systems play an essential role in limiting prion-like propagation by degrading seeding species, constraining trans-conformational changes, and removing protein aggregates through proteolytic pathways, thereby preserving cellular homeostasis. The major proteolytic routes include the ubiquitin-proteasome system (UPS), autophagy, and the endolysosomal system, all of which rely on key regulatory hubs that direct substrates to their respective degradative machineries. Among these regulators, VCP/p97 plays a central role in routing protein substrates to the proteasome, autophagy, and lysosomes. It participates in numerous degradative processes, including macroautophagy, mitophagy, the unfolded protein response (UPR), UPS function, endolysosomal activity, ER-associated degradation (ERAD), and translational control (Meyer and Weihl, 2014). Its pleiotropic contribution to proteostasis is enabled by the VCP homohexamer’s interaction with a growing set of adaptors and cofactors, which guide the processing of cytosolic and membrane-associated substrate proteins.

Mutations in the VCP gene are associated with frontotemporal lobar degeneration and other neurodegenerative disorders. P97/VCP is a homohexameric AAA-ATPase complex that primarily functions as an unfoldase. VCP has been implicated in several aspects of the Tau prion-like process, including the amplification of Tau seeding (Batra et al., 2024), the clearance of Tau aggregates (Giong et al., 2024), and the removal of damaged endolysosomes induced by Tau fibrils (Papadopoulos et al., 2017). VCP has also been suggested to modulate amyloid precursor protein (APP) metabolism through various mechanisms, including a selenoprotein-dependent pathway (Jang et al., 2017), the regulation of extracellular vesicle secretion (Lu et al., 2025), and its involvement in the secretory pathway, where it mitigates the toxic effects of ectopically expressed APP (Yu et al., 2024).

APP is a type I transmembrane protein that is processed to generate Aβ peptides, which are the main constituents of amyloid deposits in AD. The neuronal isoform APP695 is translated in the endoplasmic reticulum (ER), and the mature glycosylated protein is trafficked to the plasma membrane via the secretory pathway (Suh and Checler, 2002). APP is cleaved by the α-secretase ADAM10 within the Aβ region, releasing the extracellular fragment sAPPα and the membrane-bound C-terminal fragment αCTF (C83). This cleavage prevents Aβ formation. In the endocytic pathway, APP is processed by the β-secretase BACE1, producing βCTF (C99), the direct precursor of Aβ peptides. Both αCTF and βCTF are subsequently cleaved by γ-secretase (Presenilin 1), generating Aβ or the p3 fragment, along with the APP intracellular domain (AICD) (Vrancx and Annaert, 2025). Within the ER, a ribosome-associated quality control system regulates the levels of ectopically expressed APP, relying on IRE1, the E3 ubiquitin ligase Hrd1, and p97/VCP (Li et al., 2023). Another ER-associated protein, Seleno S, facilitates the transfer of misfolded proteins from the ER to the cytosol and is required for the degradation of βCTF/C99, thereby reducing Aβ production—a mechanism also shown to depend on the VCP–C99 interaction (Jiang et al., 2017). More recently, VCP has been suggested to regulate APP processing along the endolysosomal pathway; VCP inhibition increases the release of APP metabolites and Aβ peptides within extracellular vesicles originating from late endosomal multivesicular bodies (Lu et al., 2025; Vingtdeux et al., 2007; Rajendran et al., 2006).

Together, these findings highlight the potential contribution of the VCP AAA-ATPase to key pathogenic mechanisms underlying neurodegenerative diseases, including AD. By regulating ERAD, autophagy, and vesicular trafficking, p97/VCP may influence APP metabolism, but the precise cellular and molecular mechanisms remain unclear. In this study, we investigate how VCP interacts with APP to regulate its trafficking and secretase-dependent processing, thereby affecting the stability of full-length APP and the production of Aβ peptides.

## 1. Materials and methods

### 1.1. Antibodies and reagents

APP-Cter-C17 (1/4,000) is a homemade rabbit antiserum directed against the last 17 amino acids of APP (Sergeant et al. 2002; Vingtdeux et al. 2007). Mouse anti-β-actin (1/10,000) A5441 was obtained from Sigma. Rabbit anti-calnexin (1/2,000) 2679S and mouse anti-p62/SQSTM1 (1/2,000) ab56416 antibodies were purchased from Abcam and Cell Signaling Technology, respectively. Mouse anti-poly-ubiquitinylated conjugates (1/1,000) BML-PW8805-0500 were obtained from Enzo Life Sciences. Mouse anti-VCP (1/2,000) MA3-004 was purchased from Invitrogen. Secondary antibodies (horseradish peroxidase-labeled goat anti-rabbit IgG, 1/5,000 or horseradish peroxidase-labeled horse anti-mouse IgG, 1/50,000) were obtained from Vector Laboratories. The horseradish peroxidase-conjugated goat antibody specific for µ-chain was purchased from Sigma-Aldrich. NMS-873, the allosteric and specific VCP ATPase inhibitor, was purchased from Sigma-Aldrich and was used at the indicated concentrations. Cycloheximide (CHX) was purchased from Tocris Bioscience.

### 1.2. Cell culture

SKNSH-SY5Y (SY5Y) cells stably expressing the human APP695 isoform (SY5Y-APP695) were obtained through lentiviral transduction of the human APP 695 cDNA under the PGK promoter. Clone selection was carried out by limiting dilution of SY5Y transduced cells at 0.5 cells per well in 96-well plates. Clones were expanded for 2 to 3 weeks before selection, as confirmed by Western blotting and APP695 expression. Selection was based on APP695 expression levels similar to those in the human brain. Additionally, SH-SY5Y cells inducibly expressing the C99 fragment (SY5Y-C99) were previously described (Becot et al., 2020; Lauritzen et al., 2019). Briefly, SY5Y cells stably expressing the non-mutated human neuronal APP695 isoform or the C99/CTF APP fragment were maintained in Dulbecco’s Modified Eagle Medium (GIBCO by Life Technologies) supplemented with 5% fetal bovine serum, 2 mM L-glutamine, 1 mM non-essential amino acids, and penicillin/streptomycin (GIBCO by Life Technologies). These cells were cultured at 37°C in a 5% CO_2_ atmosphere with G418 or puromycin, respectively, to generate stable lines. SY5Y-C99 expression was induced in SY5Y-C99 cells by the addition of doxycyclin (10µg/mL).

### 1.3. siRNA transfection and ΔENotch transfection

One day before transfection, cells were plated into 12-well plates. Transient transfections of cells were carried out using ON-TARGETplus Non-targeting Pool (siRNA Negative Control), the ON-TARGETplus Human VCP (7415) siRNA obtained from Dharmacon, or SQSTM1/p62 siRNA (ThermoFisher) at 10 pM final using Lipofectamine^TM^ RNAi MAX (Invitrogen) according to the manufacturer’s instructions. After 72 h, transfected cells were treated with drugs or directly collected for further analysis. For ΔENotch, cells were plated in 12-well culture dishes at a density of 250,000 cells per well and transfected 24 h later with Lipofectamine 2000 (Invitrogen) according to the manufacturer’s instructions, using 1 µg of ΔENotch plasmid as previously described (Sergeant et al., 2019). After 72 h, transfected cells were treated with drugs or directly collected for further analysis.

### 1.4. Drug treatments

Cells were plated into 12-well plates and cultured for one day before drug treatment. The following day, the cell medium was replaced with fresh medium containing drugs at specified concentrations, and cells were exposed for the designated durations. After treatment, the culture medium was collected and stored at -80°C until use to measure Aβ_1-40_ and Aβ_1-42_ peptide concentration by ELISA. Cells were rinsed once with phosphate-buffered saline (PBS) and then lysed in RIPA buffer (50 mM Tris-HCl, pH 7.4, 100 mM NaCl, 10 mM MgCl_2_, 5% NP-40, 20% glycerol, and 0.5% sodium dodecyl sulfate), supplemented with protease inhibitors and benzonase. Total protein concentration was determined using the Pierce BCA Protein Assay Kit (Thermo Scientific) according to the manufacturer’s instructions.

### 1.5. Cell surface biotinylation

SY5Y-APP695 cells were plated at a density of 5.91 x 10^4^ cells into 100 mm^2^ dishes and cultured with 10 mL of supplemented DMEM cell medium 24 hours before siRNA transfection or at a density of 9.81 x 10^4^ cells before drug exposure. At the end of treatments, cell-surface proteins were biotinylated using Sulfo-NHS-SS-Biotin, as recommended by the manufacturer, using the Pierce Cell Surface Protein Isolation Kit (Thermo Scientific). Briefly, cells were incubated for 30 minutes at 4°C on a rocking platform with Sulfo-NHS-SS-Biotin. Cells were lysed, and labeled proteins were isolated with NeutrAvidin agarose beads. Neutravidin-binding biotinylated proteins were eluted from the beads with NuPAGE® lithium dodecyl sulfate (LDS) 2X lysis buffer added with 50 mM dithiothreitol, heated for 5 minutes at 95°C, and further analyzed by western blot.

### 1.6. VCP interactome

SY5Y-APP695 cells were cultured in DMEM supplemented with 10% FBS, 2 mM L-glutamine, 1 mM NEAA, and penicillin-streptomycin (GIBCO by Life Technologies). When they reached 80% confluence, 5 µM NMS-873 was added for a 6-hour treatment. The cells were then harvested by trypsinization, washed by centrifugation (600 x g, 5 min, RT) with PBS, and resuspended in PBS. A 1 mM DSP crosslinker (Tebubio) was added for 30 minutes at room temperature. The reaction was quenched by adding 50 mM Tris-HCl at pH 7.5 (final concentration) for an additional 10 minutes at room temperature. Cells were then pelleted by centrifugation and lysed with lysis buffer [25 mM Tris pH 7.5, 150 mM NaCl, 1% NP-40, 1 mM EDTA, 5 mM MgCl_₂_, 0.5% Na deoxycholate, and complete protease inhibitors (Roche)], followed by mild sonication on ice. Proteins were recovered from the lysate by centrifugation (21,000 × g, 30 min, 4°C). The supernatant was transferred to a clean tube and subjected to the BCA protein quantification assay (Thermo) to determine the total lysate protein concentration. Immuno-affinity recovery of the VCP protein complex was performed by incubating the sample with (i) 0.7 µg/mg protein of VCP antibody (MA3-004, Thermo Fisher) or Isotype IgG1k mouse control (BD Biosciences 550617) overnight at 4°C, and (ii) 50 µL/mg protein A/G magnetic bead suspension (Pure Proteome-Merck) for an additional 2 hours at 4°C. To elute immunopurified protein complexes, the beads pellet was washed 3 times 5 min with 500 µL of lysis buffer and finally resuspended in 30µL 2X SDS reducing Sample Loading Buffer (125mM Tris-HCl (pH 6.8), 2% (w/v) SDS, 20% glycerol, 100mM DTT, 0.01% (w/v) bromophenol blue) and incubated at 95°C for 5 minutes. The total 30 µL volume of immunopurified complexes was resolved by SDS-PAGE, and the acrylamide gel was stained with Colloïdal Gold Coomassie Blue. Coomassie blue-stained protein bands were processed for mass spectrometry analysis.

### 1.7 Proteomics mass spectrometry analysis

In-gel tryptic digestion was performed as previously described (Miguet et al., 2009). The UltiMate 3000 RSLCnano System (Thermo Fisher Scientific) was used to separate digested peptides. Peptides were automatically fractionated using a C18 reversed-phase column (75 µm × 250 mm, 2 µm particles, PepMap100 RSLC column, Thermo Fisher Scientific, at 35 °C). Trapping was performed for 4 minutes at 5 µL/min with solvent A (98 % H₂O, 2 % ACN, 0.1 % FA). Elution was achieved using two solvents, A (0,1 % FA in water) and B (0,1 % FA in ACN), at a flow rate of 300 nL/min. Gradient separation involved 3 minutes at 3% B, 110 minutes from 3% B to 20% B, 10 minutes from 20% B to 80% B, and the last 15 minutes at 80% B. The column was equilibrated for 6 minutes with 3% buffer B before analyzing the next sample.

The eluted peptides from the C18 column were analyzed using Q-Exactive instrument (Thermo Fisher Scientific). The electrospray voltage was set to 1.9 kV, and the capillary temperature was 275 °C. Full MS scans were collected in the Orbitrap mass analyzer over an m/z range of 400–1200 with a resolution of 70,000 at m/z 200. The target value was 3.00E+06. The fifteen most intense peaks with charge states 2-5 were fragmented in the HCD collision cell with a normalized collision energy of 27%, and tandem mass spectra were acquired in the Orbitrap mass analyzer at a resolution of 17,500 at m/z 200. The target value for this was 1.00E+05. The ion selection threshold was 5.0E+04 counts, with maximum ion accumulation times of 250 ms for full MS scans and 100 ms for tandem mass spectra. Dynamic exclusion was set to 30 seconds.

Raw data collected from nanoLC-MS/MS analyses were processed and converted into *.mgf peak list format using Proteome Discoverer 1.4 (Thermo Fisher Scientific). MS/MS data was analyzed with the search engine Mascot (version 2.4.0, Matrix Science, London, UK) installed on a local server. Searches were carried out with a mass tolerance of 0.2 Da for precursor ions and 0.2 Da for fragment ions, against a combined target-decoy database built from the human SwissProt database (taxo 9606, February 2016, 20173 entries) fused with a list of common contaminants (119 entries). Cysteine carbamidomethylation, methionine oxidation, protein N-terminal acetylation, and cysteine propionamidation were included as variable modifications. Up to one missed trypsin cleavage was allowed. For each sample, peptides were filtered using Scaffold 4.4.0 based on criteria including at least one peptide longer than 9 residues, ion score > 25, and an identity score > 0 to ensure a 1% false positive rate. Spectral counting was performed with Scaffold 4.4.0; the number of spectra per protein was normalized to the spectra for VCP in the control 1 sample. Spectra with a zero value were assigned a value of 0.1, and the data for all NMS-873-treated samples were expressed as a percentage relative to the control, which is set at 100% (Supplementary Table 1).

### 1.8. Co-immunoprecipitation assays

SY5Y-APP695 or SY5Y-C99 cells from a single T75 flask were washed with PBS and resuspended in ice-cold lysis buffer (10 mM HEPES, pH 7.4; 140 mM NaCl; 0.5% NP-40; 1 tablet of complete protease inhibitor cocktail, Roche). After centrifugation at 16,000 x g for 10 minutes at 4°C, the cell lysates were precleared with Pierce Protein A/G Plus Agarose (Thermo Scientific) for 2 hours at 4°C on a rocking platform. Cell lysates were then centrifuged for 30 seconds at 1,500 x g to remove beads. Total protein concentration was determined using the Pierce BCA Protein Assay Kit (Thermo Scientific) according to the manufacturer’s instructions. One milligram of total protein was incubated overnight at 4°C on a rocking platform with the indicated antibodies. The next day, protein-antibody complexes were affinity-captured with Pierce Protein A/G Plus Agarose (Thermo Scientific) for two hours at 4°C. Immunoaffinity protein complexes were washed three times with lysis buffer and re-suspended in 80 µL of loading buffer [NuPAGE® lithium dodecyl sulfate (LDS) 2X sample buffer supplemented with 20% NuPAGE® sample reducing agents (Invitrogen)] before SDS-PAGE.

### 1.9. Immunofluorescence

Cells were plated at 3 × 10^^5^ cells per well on poly-D-lysine-coated glass coverslips. The following day, cells were fixed with 4% paraformaldehyde in PBS for 20 minutes and then rinsed three times with PBS. Then, we used a permeabilization and blocking solution (0.1% Triton X-100, 10% SVF, or 1% BSA in PBS) for 45 minutes. After three washes with PBS + 0.1% BSA (PBS-B), cells were incubated overnight at 4 °C with primary antibodies against APP-Cter-C17 (1:4,000) and VCP (1:500). The next day, coverslips were rinsed with PBS-B and incubated with anti-IgG secondary antibodies conjugated to Alexa Fluor 488 and 568 (1:1000) (Invitrogen) in PBS-B. Nuclei were visualized using DAPI (Thermo Scientific). Images were acquired using a Zeiss Axio Imager Z2 microscope (Carl Zeiss, Germany) equipped with a Hamamatsu ORCA-Flash4.0 digital camera (Hamamatsu Photonics, Japan) and an ApoTome.2 system (Carl Zeiss, Germany). For Apotome image acquisition, the Axio Imager was used with an optical plan-apochromat 63x/1.4 oil-immersion M27 objective.

Brain-free floating sections of the hippocampus from one control and three Braak stage V/VI Alzheimer’s disease patients or from 6-month-old APPxPS1 mice (40 µm thick) were washed with PBS containing 0.2% Triton X-100 (Sigma). Sections were incubated with normal goat serum for human sections (1:100; Vector Laboratories) or with « Mouse on Mouse » blocking reagent for mouse sections (1:100; Vector Laboratories) for one hour at room temperature, then washed three times. The brain sections were then incubated overnight at 4°C with primary antibodies anti-APP C-ter (APPC17, homemade, 1:500, rabbit polyclonal) and anti-VCP (Invitrogen MA3-004, 1/500, mouse monoclonal). After three washes, sections were incubated with secondary antibodies anti-mouse Alexa Fluor 488 (1:500, Life Technologies) and anti-rabbit Alexa Fluor 568 (1:500, Life Technologies). Sections were mounted on Superfrost-coated slides and treated with Soudan Black (Millipore) in 70% ethanol for 5 minutes to reduce autofluorescence. Slices were coverslipped with Vectashield DAPI mounting medium (Vector Laboratories). Images were captured using a Zeiss Axioplan Axiophot 2 Fluorescence Microscope.

### 1.10. Western-blot analysis

Cell homogenates were separated on 4-12% Bis-Tris SDS-PAGE (Invitrogen) or 16% Tris-Tricine gels (Bio-Rad) for APP-CTFs analysis. Proteins were transferred to either a 0.45 µm pore size nitrocellulose membrane (G&E Healthcare) or a 0.22 µm pore size nitrocellulose membrane for APP-CTFs analysis. Protein transfer and quality were assessed using reversible Ponceau Red staining (0.2% Xylidine Ponceau Red in 3% Trichloroacetic acid). Ponceau Red staining was removed with 25 mM Tris-HCl, pH 8.0, 150 mM NaCl, 0.1% Tween-20 (v/v) (TNT-T), and then blocked with TNT-T containing either 5% (w/v) skimmed milk (TNT-M) or 5% (w/v) bovine serum albumin (TNT-BSA). The membranes were incubated with primary antibodies against TNT-M overnight at 4°C. Following primary antibody incubation, immunocomplexes were detected with HRP-conjugated secondary antibodies. Signal intensities of protein bands were quantified using ImageQuant TL v10.2 software.

### 1.11. Quantification of Aβ _1-40/1-42_ by ELISA

Conditioned media collected after cell treatments were centrifuged at 1000 g for 5 minutes to remove cell debris. Aβ_1-40_ and Aβ_1-42_ peptide concentrations were measured using Aβ_1-40_ and Aβ_1-42_ Human ELISA kits (Invitrogen) following the manufacturer’s instructions.

### 1.12. Statistical Analysis

The number of experiments per condition is shown in the figure legends. Statistical analyses were conducted using GraphPad Prism 10.2 software, employing the unpaired Student’s t-test for pairwise comparisons and one-way ANOVA with Bonferroni’s multiple comparisons test for multiple pairwise comparisons. Statistical significance was defined as ∗p < 0.05, ∗∗p < 0.01, ∗∗∗p < 0.001, and ∗∗∗∗p < 0.0001. All data are presented as mean ± standard error of the mean (SEM) from at least n = 3 experiments.

## 2. Results

### 2.1 VCP colocalizes and interacts with APP

VCP and APP expression were assessed in brain tissues from control subjects and individuals with Alzheimer’s disease (AD), as well as in APPxPS1 transgenic animals (Fig. 1A-C). VCP staining appeared in the cell bodies of brain tissue and co-labeled at discrete puncta with APP, with no clear differences between control and AD human brains (Fig. 1A-B). This cytosolic co-labeling was also observed in the brain tissue of APPxPS1 transgenic animals (Fig. 1C). In S5Y5-APP695 cells stably expressing the neuronal isoform APP695 (SY5Y-APP695), APP and VCP labeling partially overlapped (Fig. 1D), indicating that VCP and APP may associate at specific cellular sites and potentially form part of a shared protein complex. To explore this possible interaction, VCP was shown to co-immunoprecipitate after APP immunocapture, and vice versa (Fig. 1E). Both immature and mature APP proteins co-immunoprecipitated in S5Y5-APP695 cells transiently expressing the VCP-V5 construct (Fig. 1E). This interaction was further validated by isolating VCP immunocomplexes and analyzing the VCP interactome via mass spectrometry (Fig. 1F and Fig. 2). Over 2,349 polypeptides were identified within the immuno-isolated VCP and crosslinked protein complex (Supplementary Table 1), including 494 proteins previously reported in other VCP interactome databases as VCP adaptors, co-factors, or VCP ATPase substrates (Fig. 1F). APP and several proteins associated with APP were detected within the VCP interactome (Fig. 2A). STRING-db analysis (Szklarczyk et al., 2023 and 2025) of the VCP interactome suggests an indirect interaction between VCP and both immature and mature full-length APP proteins, implying that VCP’s regulatory role in APP metabolism may involve either APP, VCP, or their respective co-factors. Proteins identified as part of the shared VCP/APP interactome were mainly related to catabolic or proteolytic regulation, with a lesser connection to autophagy (Fig. 2B).

**Figure 1:**
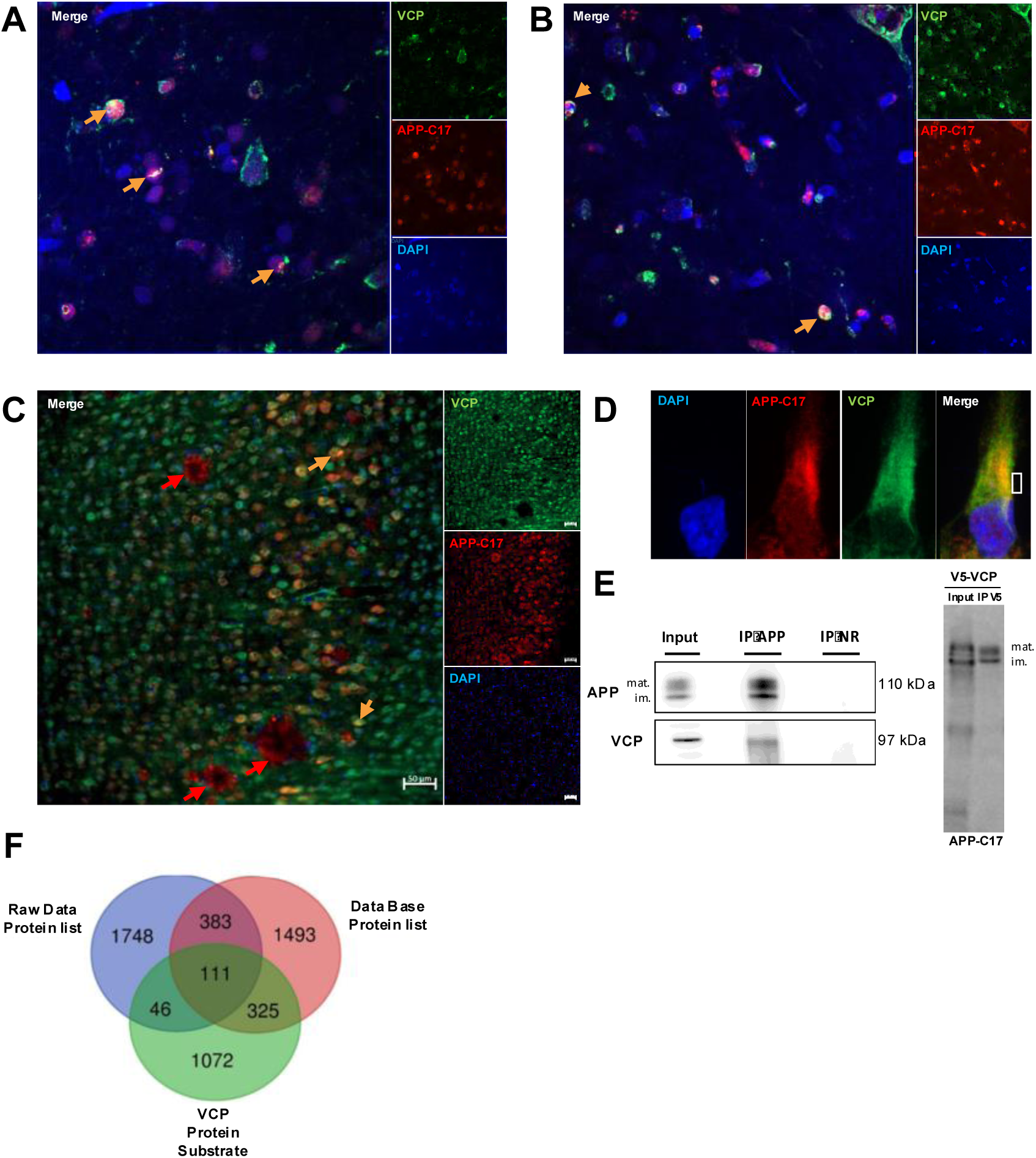
VCP and APP colocalization and interaction. (A and B) Post-mortem human cortical brain tissues from one control (A) and three Alzheimer’s disease patients (B) were immunostained with VCP (Green) and APP (Red). Cell nuclei are stained in blue using DAPI. Orange arrows indicate areas of VCP and APP overlap staining. (C) In the APP/PS1 murine transgenic model, VCP co-labels neurons that accumulate APP metabolites (APP-CTFs, shown by the orange arrows). Amyloid plaques are not labelled by the VCP antibody (indicated by the red arrows). (D) VCP colocalizes with APP in SY5Y-APP695 cells, as shown by VCP and APP co-labelling. (E) Immunoprecipitation of APP using antibodies targeting APP-Cter-C17 and VCP staining, as well as irrelevant immunoglobulin (IP NR) as a control, indicates an interaction between APP and VCP in the same APP immunopurified complex. Immunoprecipitation of V5-VCP following transfection of SY5Y-APP695 pull-down APP immature (im.) and mature (mat.) proteoforms (E, right panel) (D) Venn diagram of VCP interactome proteins (2288) identified by mass spectrometry MS/MS in the present study, with the identified VCP interactors from public databases and referenced VCP protein substrates.

**Figure 2:**
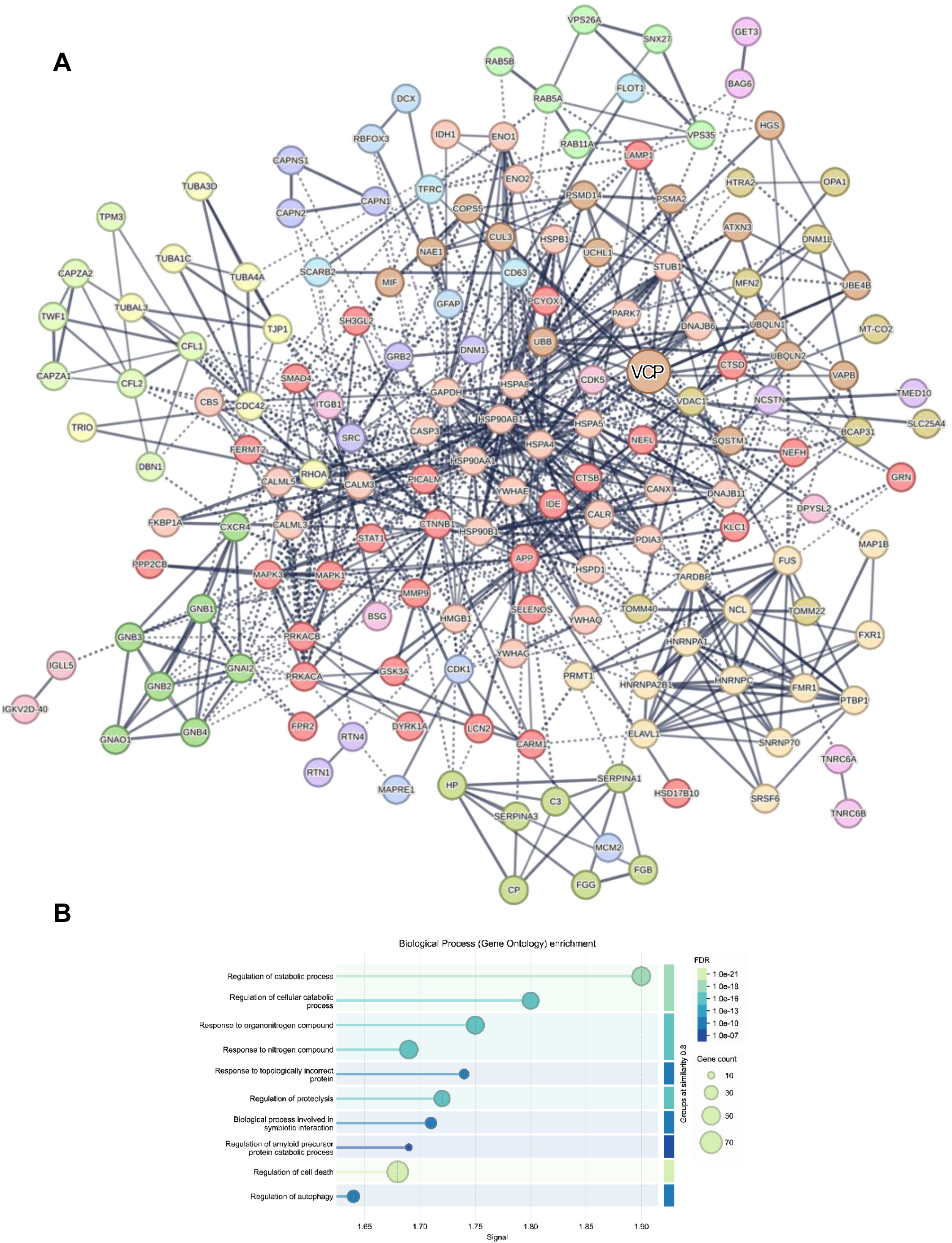
VCP and APP protein interaction network modulated by VCP ATPase inhibitor. (A) VCP /APP interactome proteins that were modulated following VCP ATPase inhibition with NMS-873 were processed using STRING (www.string-dg.org) to generate a protein-protein interaction network of 180 VCP and APP proteins interacting and modulated by NMS-873 treatment of S5Y5-APP695, utilizing a high-confidence (0.7) interaction score (Szklarczyk et al., 2023 and 2025). (B) STRING gene ontology analysis was used to identify the major biological processes of VCP and APP interactors, highlighting the regulation of catabolic processes, including proteolysis regulation, APP catabolic process, and autophagy as the primary biological processes associated with the VCP / APP interactome.

### 2.2 VCP ATPase activity regulates APP metabolism and Aβ production

The potential modulation of APP metabolism was initially studied by silencing VCP expression with siRNA to mimic a loss of VCP function, which is relevant to the loss of VCP observed in neurodegenerative diseases (Fig. 3) (Darwish et al., 2020; Wani et al., 2020; Chu et al., 2023). We evaluated APP and its metabolites produced from both the amyloidogenic and non-amyloidogenic pathways. Four small VCP-interfering RNAs (siRNAs) were combined into a pool and transfected into SY5Y cells stably expressing human SY5Y-APP695. The effects of VCP silencing were assessed 72 hours after transfection (Fig. 3). After VCP silencing, VCP levels decreased by 75% compared to the original endogenous expression. As expected, VCP silencing caused an accumulation of poly-ubiquitinylated proteins, indicating a loss of VCP function (Fig. 3A and 3B). In fact, VCP silencing results in the buildup of VCP client proteins, mainly poly-ubiquitinylated proteins targeted for degradation by the 26S ubiquitin-proteasome system (Li et al., 2024). Next, we examined how VCP downregulation affected APP maturation and processing. First, we analyzed full-length APP expression. Full-length APP695 appears as two bands at 110 kDa and 115 kDa (Fig. 3A), representing the immature or neosynthesized form and the glycosylated mature form, respectively. VCP silencing caused a significant increase in immature APP levels compared to control conditions and, to a lesser extent, in mature APP levels (Fig. 3A and 3C) (Fig. 3A and 3C).

**Figure 3:**
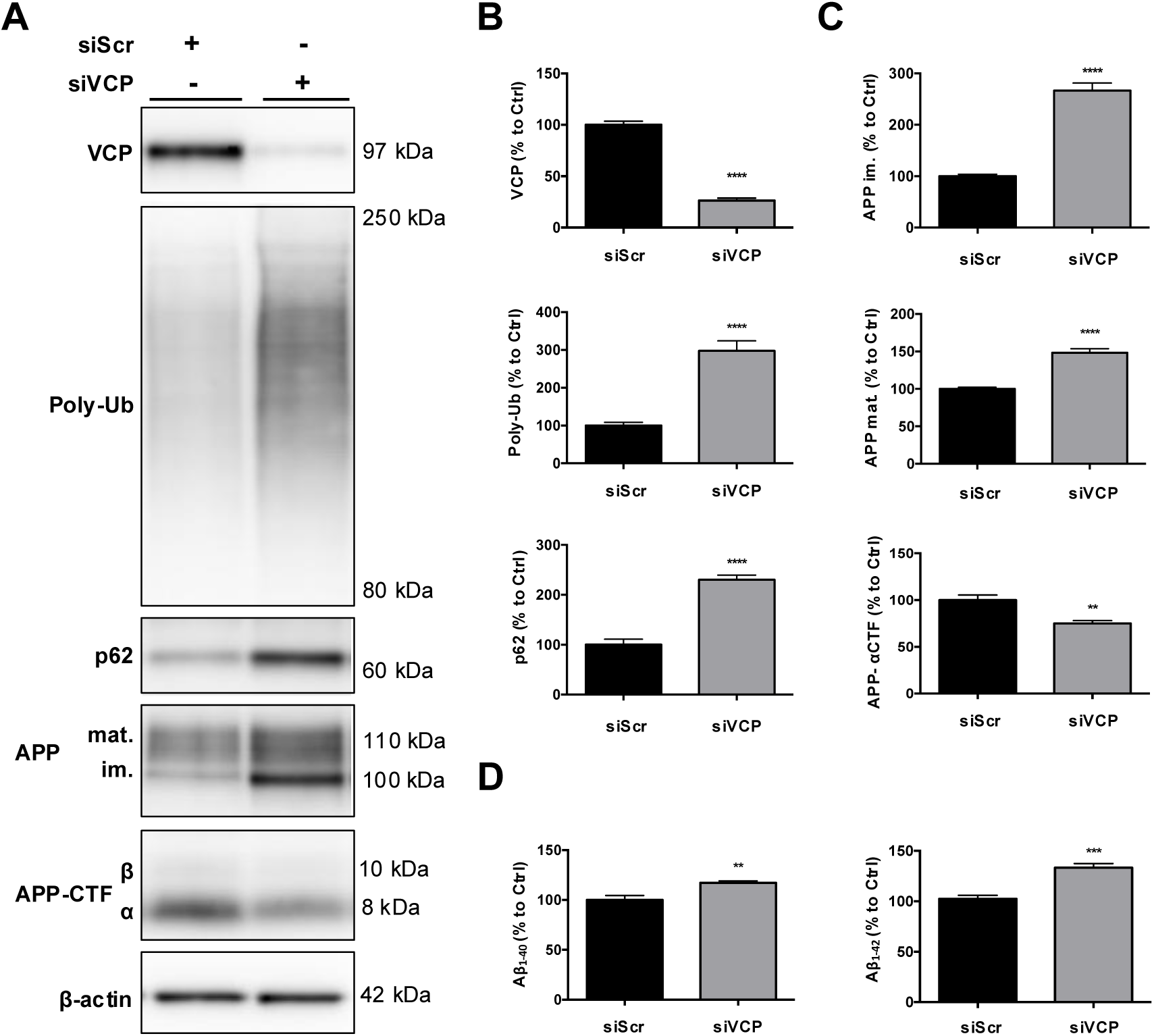
Effect of VCP loss of expression on APP metabolism in SY5Y-APP695 cells. (A) SY5Y-APP695 cells were transfected with scrambled siRNA (siScr) or VCP siRNA (siVCP) for 72 hours. (A) SDS-PAGE resolved cell protein lysates, and transferred membranes were blotted with antibodies targeting VCP, poly-ubiquitinated proteins (Poly-Ub), SQSTM1 (p62), APP-Cter-C17 (APP), and β-actin as a protein loading control. Apparent molecular weights are indicated to the left of the Western blots in kilodaltons (kDa) (B and C). Densitometric analyses and semi-quantification of VCP, Poly-Ub proteins, immature (mi.) and mature (mat.) APP levels, and APP α-carboxy-terminal fragments (APP-αCTF). (D) The concentrations of Aβ_1-40_ and Aβ_1-42_ peptides were measured in the cell-conditioned media by ELISA. Histograms represent the mean ± SD on the y-axis and are expressed as a percentage of the control value, arbitrarily set at 100%. n=6 *p<0.05, **p<0.01, ***p<0.005, ****p<0.001 Unpaired t-test.

Full-length APP is cleaved by the α- or β-secretase to produce soluble APP (sAPP), and APP-carboxy-terminal fragments αCTF/C83 or βCTF/C99, respectively. The γ-secretase then processes these APP-CTFs to release the APP intracellular domain (AICD) and the p3 or Aβ peptide from the α- or βCTF, respectively. Silencing VCP significantly reduced APP-CTF levels (Fig. 3A). Specifically, αCTF expression decreased by 25% (Fig. 3C). βCTF could not be quantified in control and siVCP-transfected conditions; however, Western blot analysis showed a decrease in the βCTF band after VCP silencing compared to control (Fig. 3A). Additionally, the concentrations of Aβ_1-40_ and Aβ_1-42_ peptides in the cell media increased by 17% and 33%, respectively, following VCP silencing (Fig. 3D). Therefore, VCP regulates APP metabolism at two levels: it influences full-length protein expression and APP secretase processing, both affected by VCP silencing in SY5Y-APP695.

Next, we assess whether VCP protein expression or its ATPase activity affects APP metabolism. VCP ATPase activity decreases with increasing concentrations of NMS-873 (Fig. S1A), which is the most potent allosteric and specific VCP ATPase inhibitor with an IC_50_ of 365 nM, as demonstrated *in vitro* using recombinant VCP protein. In SY5Y-APP695 cells, VCP inhibition by NMS-873 caused a gradual accumulation of poly-ubiquitinylated proteins (Fig. S1A and S1D). VCP levels remained unchanged (data not shown). Like VCP silencing, VCP inhibition with NMS-873 led to a dose-dependent increase in immature APP levels (Fig. S1B). A significant decrease in APP-CTFs was observed starting at 500 nM NMS-873, with αCTFs dropping by 15-20% after treatment (Fig. S1B and S1C). Levels of Aβ_1-40_ and Aβ_1-42_ peptides in cell media increased by 20-60% and 16-40%, respectively (Fig. S1E). Therefore, inhibiting VCP ATPase activity mimics the effects of VCP protein loss, indicating that both VCP expression and ATPase activity are essential for regulating APP levels and Aβ peptide production.

### 2.3 VCP regulates the rate of trafficking of APP to the plasma membrane

To understand how VCP regulates full-length APP expression, we examined the effect of VCP silencing on APP degradation following cycloheximide treatment to block protein synthesis. VCP silencing led to the accumulation of both immature and mature APP (Fig. 4A), indicating that the increased levels of full-length APP were not due to enhanced APP translation. To measure the rate of full-length APP production, SY5Y-APP695 cells were treated with the protein synthesis inhibitor cycloheximide (CHX) for various time periods (Fig. 4A). APP levels were monitored at specific intervals to track the turnover of immature and mature full-length APP (Fig. 4A-C). In the control, APP levels began to decline 15 minutes after CHX addition. Nearly all APP was degraded within 3 hours, with an average 25% loss every 30 minutes, consistent with previous studies (Storey et al., 1999) (Fig. 4A-C). VCP silencing significantly slowed down immature-full-length APP turnover, extending the half-life of immature APP by 50% (Fig. 4B). In contrast, there was no significant difference in the half-life of mature APP between control and siVCP-transfected cells, indicating that in the absence of VCP, the immature full-length APP accumulates but is less converted into mature form (Fig.4C). This degradation pattern of mature APP indicates that VCP affects the rate of APP maturation.

**Figure 4:**
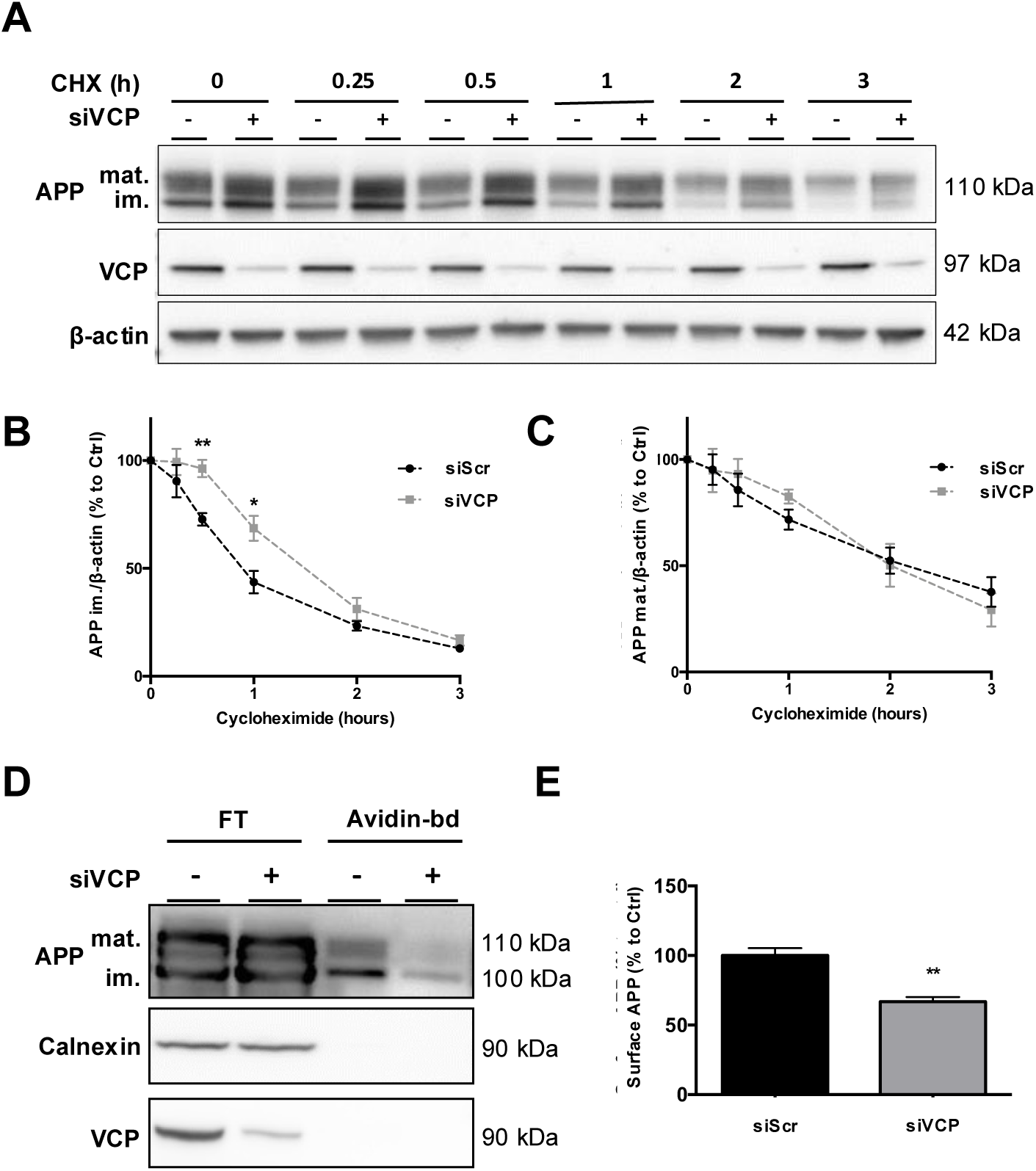
VCP loss of expression stabilizes immature and mature APP proteoforms in SY5Y-APP695 cells. (A) SY5Y-APP695 cells were transfected with negative control scrambled siRNA (siVCP -) or with VCP siRNA (siVCP +) for 72h and then treated with 40µg/mL of cycloheximide (CHX) for the indicated times from 0 to 3 hrs. Western membranes were immunoblotted with the following antibodies: APP-Cter-C17 (APP im. and APP mat.), VCP, and β-actin as a loading control. (B and C) Densitometric analyses and quantification of immature and mature APP levels were expressed as a percentage of the control (Ctrl) on the y-axis, over CHX treatment time on the x-axis. The black dotted lines represent quantification for the scrambled siRNA, while the grey dotted lines represent the siVCP condition (D). VCP siRNA reduces cell surface APP levels in SY5Y-APP695 cells. SY5Y-APP695 cells were transfected with scrambled siRNA (-) or with VCP siRNA (+) for 72h. Cell-surface proteins were chemically biotinylated and isolated with immobilized avidin beads. Flow-through and avidin-bound proteins were resolved by SDS-PAGE, and APP, VCP, and Calnexin (a cytosolic protein) were detected using antibodies against APP (APP Cter-C17), VCP, and calnexin. (E) Densitometric analysis and quantification of total APP levels. The graph indicates the mean ± SD. n=4 *p<0.05, **p<0.0, Unpaired t-test.

To explore whether VCP invalidation could affect APP transport to the cell surface, extracellular protein biotinylation experiments were performed in SY5Y-APP695 cells to measure the amount of APP addressed to the cell membrane. We found that both immature and mature APP were biotinylated at the cell surface in untransfected cells (Fig. 4D). Silencing VCP led to a significant 34% drop in both immature and mature APP levels at the cell surface (Fig. 4D-E). These findings suggest that VCP controls the rate of APP transport within the cell’s secretory pathway.

### 2.4 VCP regulates APP-CTFs degradation

Reduced αCTFs and increased Aβ levels may result from the slow kinetics of APP trafficking to the plasma membrane, due to VCP loss-of-function. However, we cannot rule out a direct modulatory effect of VCP on APP-CTFs and Aβ. Therefore, SY5Y-C99 inducible cells were used to assess whether VCP influences the fate of APP-CTFs.

Following induction, C99 fragments stained with the APP-CterC17 polyclonal antibody were visualized in large vesicles surrounding the cell nucleus (Fig. 5A). The βCTF/C99 was co-immunoprecipitated with VCP but not with αCTF/C83 (Fig. 5B). After inhibiting VCP’s ATPase activity with NMS-873, C99 fragments were significantly reduced (Fig. 5C-5D). The levels of Aβ_1-40_ and Aβ_1-42_ in the cell medium dropped by more than 50%, indicating that VCP regulates the degradation of APP-CTFs and the production of Aβ. Potential inhibition of γ-secretase was ruled out, as Notch γ-secretase processing into NICD (De Strooper et al., 1999; Vingtdeux et al., 2007) was unaffected by NMS-873 treatment (Supplementary Fig. 2A-B). In contrast, DAPT, a γ-secretase inhibitor, abolished NICD production from the ΔENotch-expressing vector, indicating that Notch is a γ-secretase substrate distinct from APP.

**Figure 5:**
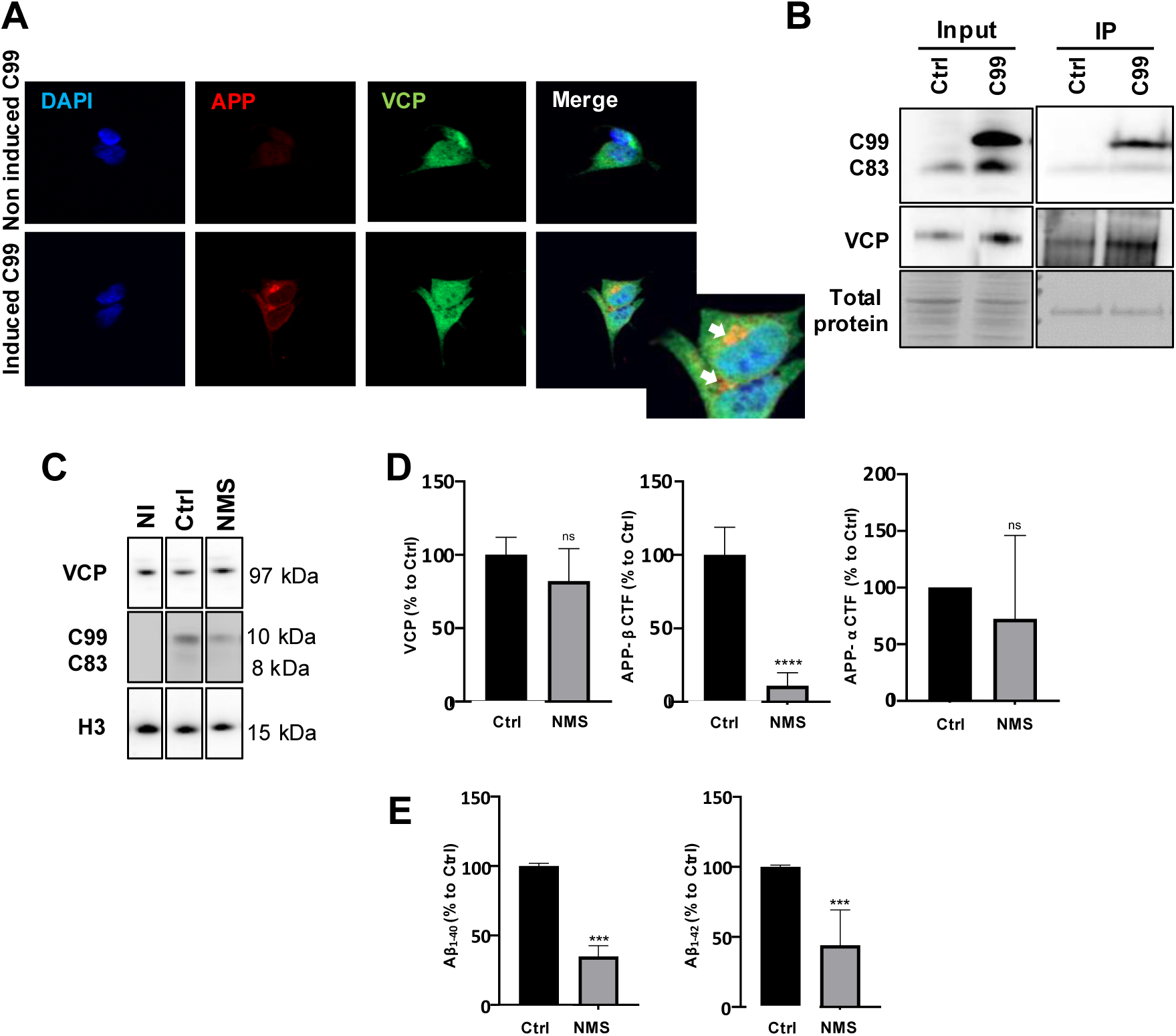
Effect of VCP ATPase inhibition by NMS-873 on C99 metabolism in inducible-C99 SY5Y human neuroblastoma cells. (A) VCP colocalizes with APP in SY5Y-C99 cells, as shown by co-labeling with APP-Cter-C17 (Red) and VCP (Green) antibodies. (B) VCP was immunoprecipitated, and C99 was detected only following C99-induced expression (C99 lane) compared to the non-induced condition (Ctrl lane). (C) SY5Y-C99 cells were treated for 24 hours with 1.5 µM of the allosteric VCP ATPase inhibitor NMS-873 as indicated. (C-D) SDS-PAGE-resolved cell protein lysates and nitrocellulose membranes were immunolabeled with the following antibodies: VCP, APP-Cter-C17, and Histone H3 as a loading control. (E) ELISA analysis of conditioned medium from SY5Y-C99 cells was performed to quantify secreted Aβ_1-40_ and Aβ_1-42_. Histograms show the mean ± SD. n=3; ns=non-significant, *p<0.05, **p<0.01, ***p<0.001. Data were analyzed using an unpaired t-test or Mann-Whitney test.

### 2.5 VCP regulation of APP metabolism and the ubiquitin-proteasome system

VCP is a homohexameric protein complex that functions as a hub, engaging with multiple adaptors and co-factors, many of which are associated with ubiquitin-proteasome or autophagy-lysosome systems to control protein degradation within cells. If the proteasome is involved in VCP-dependent regulation of APP metabolism, then co-inhibition of VCP and the proteasome should change full-length APP levels, stabilize or increase APP-CTFs, and restore Aβ peptide levels (Fig. 6).

**Figure 6:**
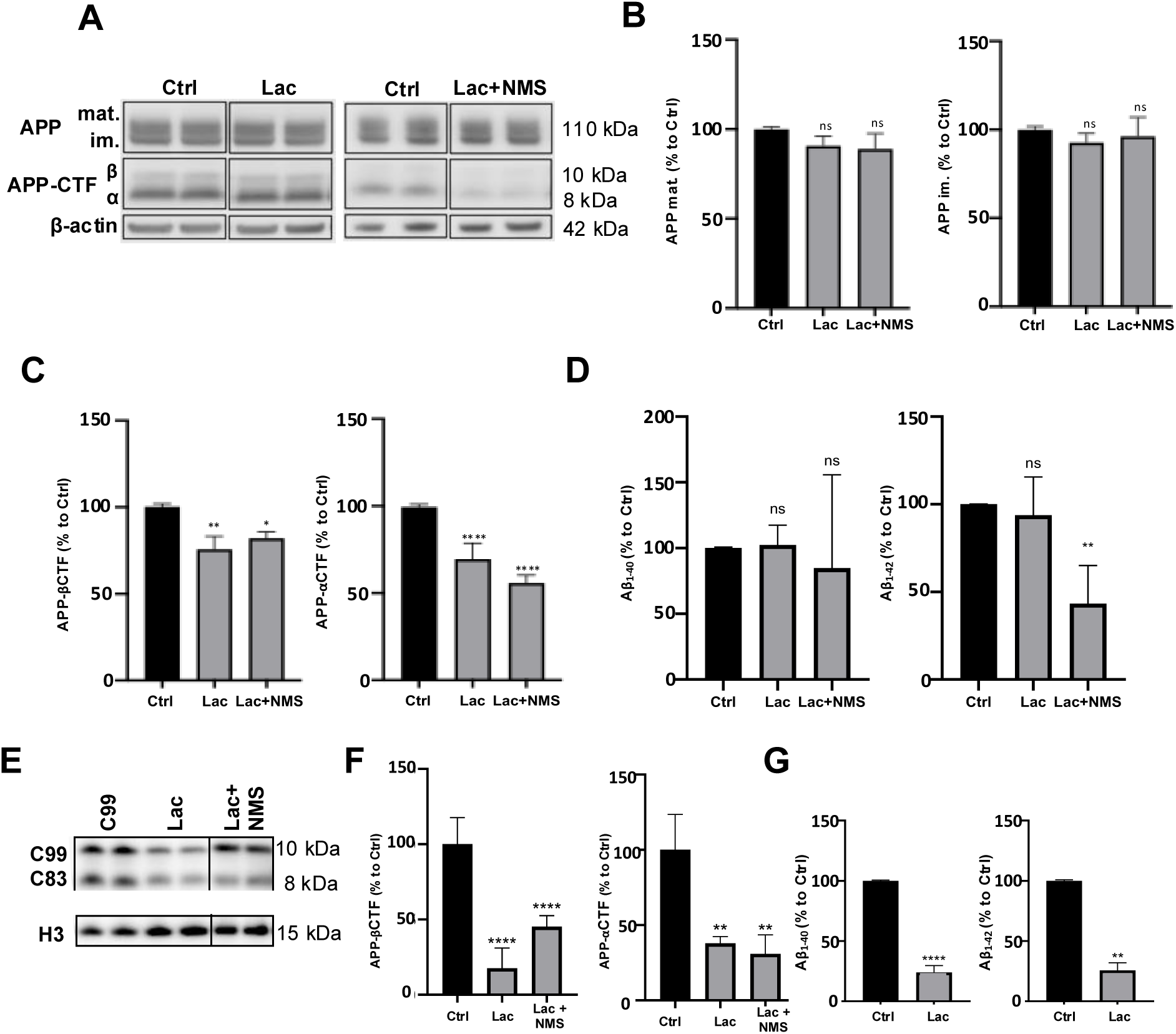
Co-inhibition with NMS-873 and Proteasome, APP, and C99 processing in SY5Y cells. SY5Y-APP695 or C99-induced SY5Y cells were treated with Lactacystin (1 µM, 24 h) alone or combined with VCP ATPase inhibitor NMS-873. After gel electrophoresis and protein transfer, Western membranes were blotted with APP-Cter-C17 and β-actin as a loading control. (A) Immunolabeling of immature (im.) or mature (mat.) proteoforms of APP, APP β- and α-CTFs at 8 and 10 kDa, and β-actin for SY5Y-APP695 cell protein lysates under control conditions (Ctrl), treated with Lactacystin (Lac), or with Lactacystin and NMS-873 (Lac+NMS). (B) Histograms representing the densitometric semi-quantification of immature (mat.) and immature (im.) APP proteoforms and (C) βCTF and αCTF, expressed on the y-axis as percentage of the control value, arbitrarily given a 100% value for the control (Ctrl: Black column). lactacystin (Lac), and lactacystin + NMS-873 (Lac+NMS) conditions correspond to the grey columns. Panels (E) to (G) are similar to panels (A) to (D) for C99-inducible SY5Y cells. C99 and C83 fragments are indicated. (D and G) ELISA quantification of secreted Aβ_1−40_ and Aβ_1−42_, expressed as a percentage of the control condition (100%). Data are shown as the mean ± SD (n = 3 independent experiments), ns = non-significant, **p < 0.01, ***p < 0.001, and ****p < 0.0001.

In SY5Y-APP695 cells, proteasome inhibition with lactacystin alone did not alter the expression levels of immature or mature APP (Fig. 6 A, B). This finding suggests that full-length APP is not degraded via the proteasome pathway. Additionally, when VCP was inhibited in the presence of lactacystin, no accumulation of full-length APP was observed (Fig. S1). These results collectively indicate that VCP’s effect on full-length APP does not involve proteasomal degradation. Co-inhibition of VCP and the proteasome produced a higher reduction for CTFs observed when only VCP was inhibited (Fig. S1). Treatment with NMS-873 induces only 10% inhibition, whereas co-inhibition of VCP and the proteasome reduces inhibition to 50% of the control level (Fig. 6C). This indicates a distinct VCP-mediated mechanism for full-length APP and APP-CTFs.

Compared to the control, Aβ_1-42_ concentration in cell media was reduced by 50% only when both VCP and the proteasome were inhibited, while Aβ peptide concentrations remained unaffected by proteasome inhibition and increased only in the presence of NMS-873 (Fig. S1). To directly assess proteasome activity on APP-CTFs, proteasome inhibition was performed in SY5Y-C99 cells. Notably, levels of C99 and C83 were significantly decreased by proteasome inhibition, VCP inhibition, and the combined inhibition of both, and levels of Aβ_1-40_ and Aβ_1-42_ dropped by up to 25% compared to control levels following lactacystin-induced proteasome inhibition. Thus, proteasome inhibition mimicked VCP inhibition in SY5Y-C99 cells. These effects were not due to changes in γ-secretase activity, as NICD production from ΔENotch remained unaffected by lactacystin or by the combined lactacystin and NMS-873 treatments (Fig. S2A-B). These results indicate that the proteasome does not mediate VCP’s modulatory effect on APP metabolism. Surprisingly, inhibiting VCP, the proteasome, or both appears to diminish APP-CTF expression.

### 2.6 VCP regulation of APP metabolism and autophagy

Protein degradation mediated by VCP can occur through autophagy. VCP indirectly regulates phagophore formation by stabilizing Beclin-1 (Hill et al., 2021; Wang et al., 2024). APP processing by secretases occurs along the endolysosomal pathway (Vingtdeux et al., 2007), which intersects with the autophagy pathway (Birgisdottir and Johansen, 2020) and is believed to contribute to the phagocytosis of APP, degradation of APP-CTFs, and Aβ through the PICALM APP interactor (Tian et al., 2013); PICALM was herein identified within the VCP interactome (Fig. 2A). PICALM was 8.9 times more abundant in the VCP interactome under NMS-873 treatment, suggesting an increase of the VCP/PICALM complexes (Supplementary Table 1).

We then assessed whether the loss of VCP activity promotes APP metabolism via an autophagy-mediated mechanism (Fig. 7A). After inducing autophagy with Torin1, an mTOR inhibitor and autophagy activator, immature APP slightly increased, while CTFs decreased, and Aβ peptides remained unchanged (Fig. 7A to 7D) under autophagy activation. Autophagy flux inhibitors chloroquine (CQ) and Bafilomycin A1 (BafA1) significantly raised the levels of both immature and mature APP (Fig. 7A and 7B). APP-CTFs were notably elevated, while Aβ_1-40_ and Aβ_1-42_ levels dropped to 20% of control levels (Fig. 7C and 7D). Aβ production was likely unaffected by γ-secretase modulation, as γ-secretase processing of ΔENotch remained unchanged with Torin1, BafA1, or CQ (Fig. S2C-D). These findings suggest that Torin1-induced autophagy activation slightly promotes APP-CTF degradation, whereas autophagy flux inhibitors cause an accumulation of APP and APP-CTFs, thereby blocking Aβ production. However, CQ and BafA1 also inhibit the endolysosomal pathway through their lysosomotropic activity (Vingdeux et al., 2007).

**Figure 7:**
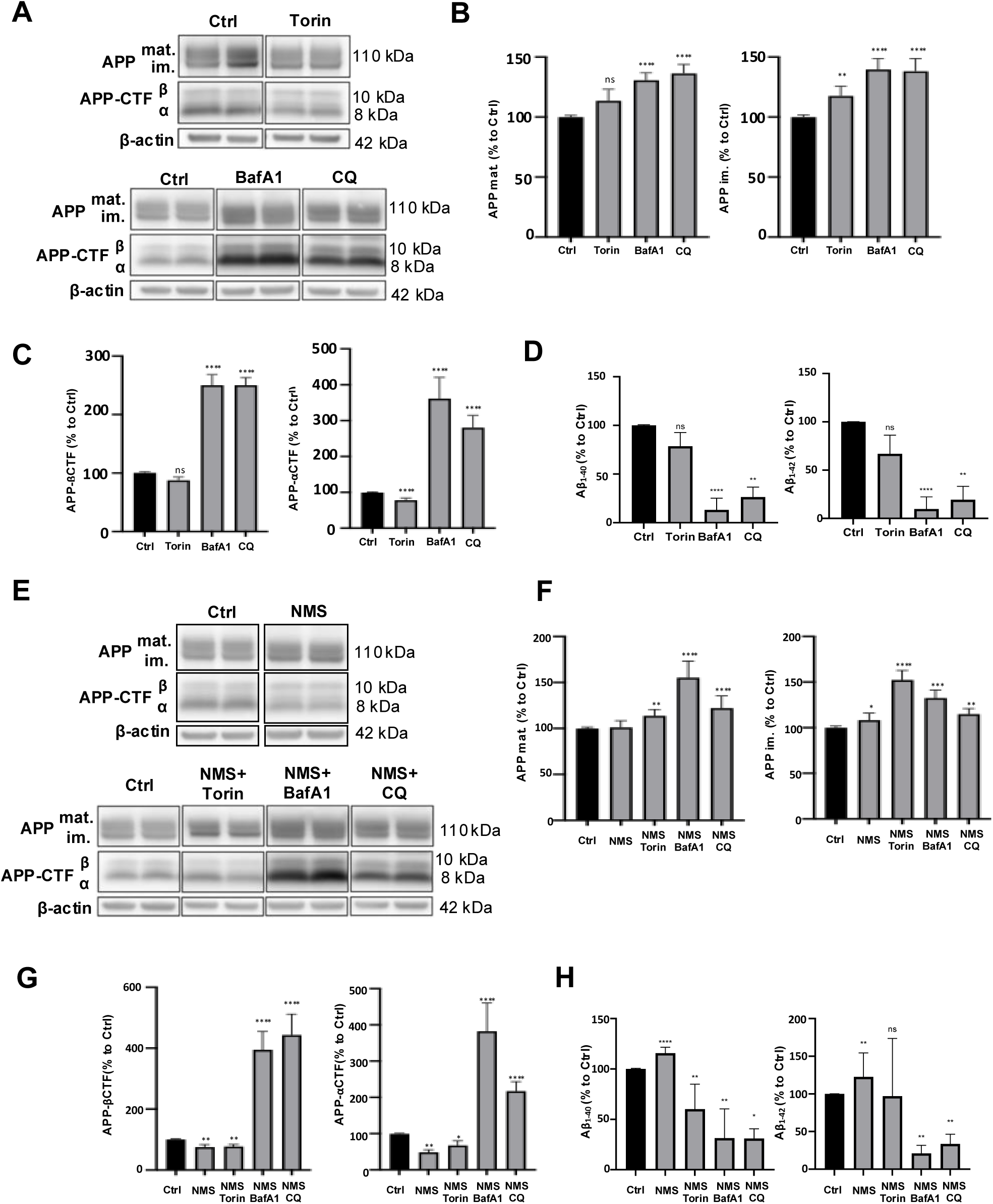
Effect of autophagy activation or inhibition without or with NMS-873 co-treatment on APP in SY5Y-APP695. (A) SY5Y-APP695 cells were treated for 24 h with autophagy activator Torin1 (1µM), autophagy flux inhibitors Bafilomycin A1 (BafA1, 100nM) or Chloroquine (CQ, 30 µM). Protein cell lysates were resolved by SDS-PAGE, transferred, and nitrocellulose membranes were blotted with APP-Cter-C17 rabbit polyclonal serum for the detection of mature (Mat.) and immature (im.) APP as well as APP carboxy-terminal fragments (APP-βCTF and APP-αCTF). β-actin staining was used as a protein loading control. Apparent molecular weight in kilodaltons (kDa) using molecular weight markers is indicated to the left of the Western blot images. Mature and immature proteoforms of APP were quantified (B) as well as α- and β-CTF (C) and Aβ_1-40_ and Aβ_1-42_ peptides (D). (E) SY5Y-APP695 cells were treated for 24 h with NMS-873 alone or NMS-873 together with the autophagy activator Torin1 (1µM), autophagy flux inhibitors Bafilomycin A1 (BafA1, 100nM) or Chloroquine (CQ, 30 µM). Following SDS-PAGE, mature and immature proteoforms of APP and APP-CTFs were detected with APP-Cter-C17 polysera and β-actin, used as a protein loading control. Immature and mature APP proteoforms were quantified (F) as well as APP-CTFs (G) and A β1-40 and 1-42 peptides (H). Data are represented as histograms of the mean ± SD (minimum of n = 3 independent experiments produced in triplicate for each experimental condition), ns = non-significant, **p < 0.01, ***p < 0.001, and ****p < 0.0001.

We then examined how autophagy affects the APP metabolism changes induced by VCP inhibition (Fig. 7E and 7F). αCTF and βCTF levels decreased after VCP ATPase inhibition (Fig. 7E-7G), and Aβ_1-40_ and Aβ_1-42_ levels increased by 120% compared to controls. The Torin1 autophagy activator did not eliminate this effect, although Aβ peptide levels were lowered. In contrast, both BafA1 and CQ fully reversed the effects of VCP inhibition. Overall, these findings suggest that VCP primarily influences the APP-CTF via the endolysosomal pathway rather than the autophagy pathway.

### 2.7 VCP regulation of C99 metabolism and autophagy

In SY5Y-C99 cells, activating autophagy with Torin1 lowered p62 levels without affecting LC3B-I expression, indicating enhanced autophagy flux (Fig. 8A-C). Both BafA1 and CQ increased p62 and the LC3BII/BI ratio, signaling inhibition of autophagy flux. Notably, the LC3B-II/B-I ratio was higher with CQ treatment (Fig. 8C), suggesting CQ’s lysomotropic activity further blocks the fusion of autophagosomes with lysosomes and prevents LC3B-II degradation in autolysosomes, while BafA1 already inhibits autophagy flux at the stage of autophagosome formation.

**Figure 8:**
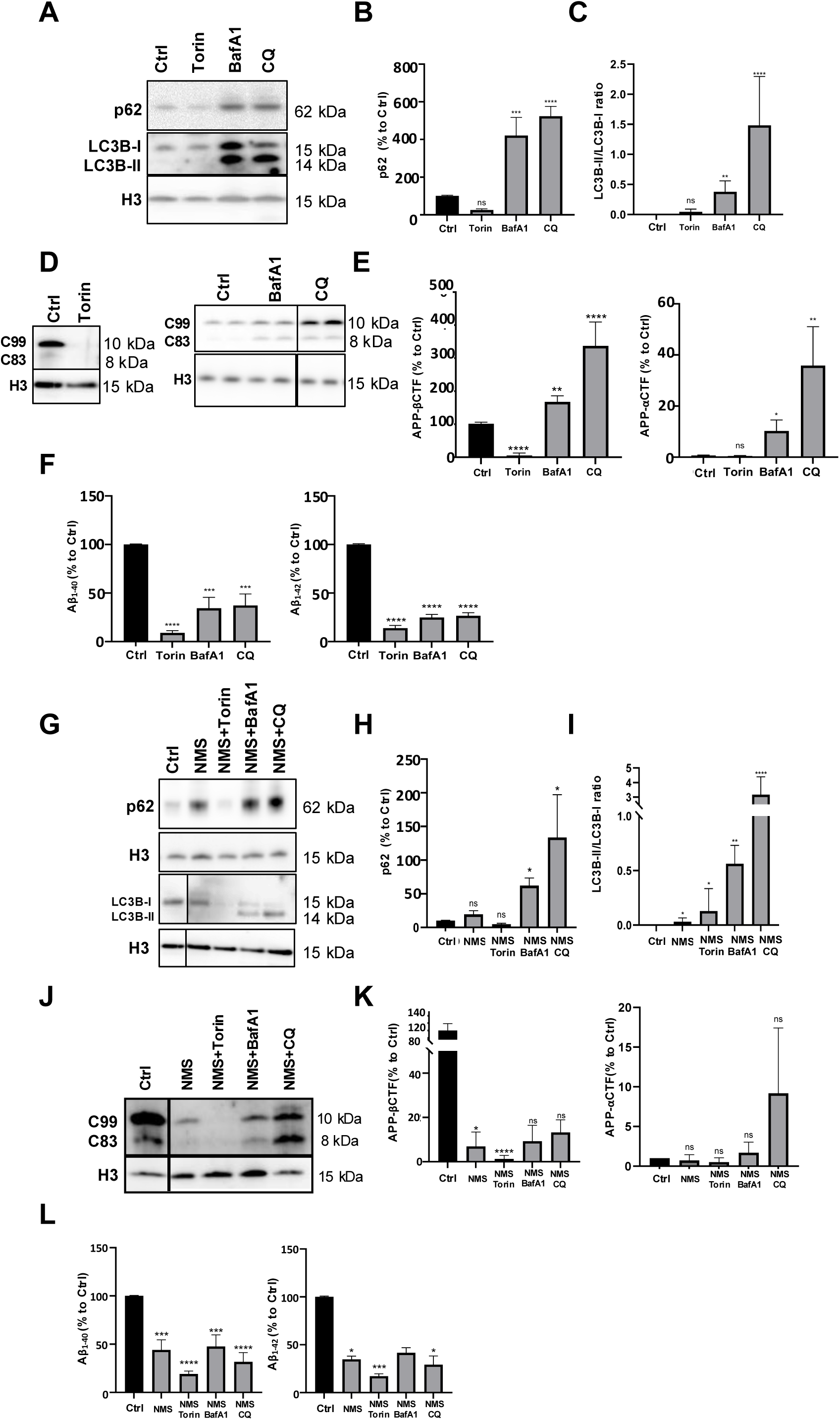
Effect of autophagy activation or inhibition with or without NMS-873 co-treatment on APP in SY5Y-APP695. (A) SY5Y cells overexpressing C99 (SY5Y-C99) were treated with the autophagy activator Torin1 (1 µM, 24 h) and autophagy inhibitors as Chloroquine (CQ, 30 µM, 24 h) and Bafilomycin A1 (BafA1, 100 nM, 24 h). (A) Cell lysates were immunolabeled with antibodies targeting p62, LC3B, and C99 and C83 fragments with the APP-Cter-C17 polysera. Histone H3 immunolabelling was used as a loading control. SQSTM1/p62 was quantified (B), and the quantitative ratio of LC3B-II / LC3B-I is represented (C). (D) C99 and C83 fragments were stained and quantified following S5Y5-C99 treatment with Torin1, BafA1, or CQ as in panel A. Quantifications of APP-CTFs and Aβ peptides are represented in panels E and F. (G to K) Following the same protocol as in panel (A) and (D) SY5Y-C99 were treated with NMS-873 together with Torin1, Bafilomycin A1 (BafA1) and chloroquine (CQ). SQSTM1/P62 and LC3B-I and LC3B-II were detected (G) following Western blotting and quantified (H and I). C99 and C83 fragments were detected with APP-Cter-C17 polysera and quantified in S5Y5-C99 protein cell lysates following co-treatment with NMS-873 and Torin1, BafA1, and CQ, panels (J and K) (L) ELISA quantification of secreted Aβ_1−40_ and Aβ_1−42_, expressed as a percentage of the control condition. Data represented as histograms of the mean ± SD (minimum of n = 3 independent experiments in triplicate for each experimental condition), ^∗^p < 0.05, ^∗∗^p < 0.01, and ^∗∗∗^p < 0.001.

Autophagy activation by Torin1 significantly reduced C99 expression and Aβ levels (Fig. 8D-8F). When autophagy flux was blocked with CQ and BafA1 (Figure 8A), levels of C99 and C83 increased substantially (Fig. 8D and E). These results demonstrate that autophagy contributes to the degradation of APP-CTFs, and when autophagy is inhibited, C99 fragments become substrates for the 𝛾-secretase to produce C83 APP-CTF fragments (to be published elsewhere). Additionally, autophagy induction reduced Aβ_1-40_ and Aβ_1-42_ production (Fig. 8F), suggesting that autophagy degrades C99 and C83, thereby preventing Aβ formation. Conversely, Aβ_1-40_ and Aβ_1-42_ were still produced under CQ and BafA1 inhibition (Fig. 8E), but their levels in the medium decreased by up to 75% compared to controls, partly due to C99 𝛾-secretase processing, which also reduces Aβ production from the 𝛾-secretase processing of C99. Gamma-secretase activity remained unchanged following BafA1 and CQ, considering the preserved 𝛾-secretase processing of Notch into NICD (Fig. S2C and S2D).

VCP ATPase inhibition with NMS-873 increased p62 expression and maintained the LC3B-II/B-I ratio, indicating an inhibitory effect on early autophagy stages (Hills et al., 2021). NMS-873 activity was counteracted by the Torin1 autophagy activator (Fig. 8G and 8H), which acts upstream of Beclin-1, whose stability is regulated by VCP. Co-treatment with NMS-873 and BafA1 or CQ further elevated p62 levels and the LC3BII/BI ratio (Fig. 8G-8I), suggesting enhanced inhibition of autophagic flux. Since VCP inhibition increased p62/SQSTM1 expression and p62 was also part of the VCP interactome, we examined the role of p62 in NMS-873 effect on APP or C99 metabolism. p62/SQSTM1 was silenced with siRNA (Fig. S3 and S4), and p62 levels were measured in SY5Y-APP695 and C99 cells under conditions of autophagy activation or inhibition, including NMS-873 treatment. In SY5Y-APP695, loss of p62 did not affect APP metabolism or C99 (Fig. S3 and S4). Co-treatment with NMS-873 and Torin1 decreased CTF levels; conversely, combined treatment with CQ and NMS-873 increased CTF levels, suggesting that C99 degradation is regulated by autophagic flux but not entirely prevented by autophagy inhibitors. This also indicates the endolysosomal pathway contributes to C99 processing, which is only partially blocked by autophagy inhibitors. The role of the endolysosomal pathway under VCP inhibition was further confirmed by immunofluorescence, showing vesicular localization of APP and APP metabolites. APP was localized to LC3B vesicles and appeared reduced following VCP inhibition by NMS-873 (Fig. 9A). As previously observed, APP was rarely localized to early endosomes and was unaffected by VCP ATPase inhibition (Fig. 9B). In contrast, after VCP inhibition with NMS-873, APP and CTFs were intensely labeled in Tsg101 late endosome vesicles (Fig. 9C), while their localization within Lamp2 lysosomal vesicles was unchanged by NMS-873 treatment. Overall, our results suggest that loss of VCP activity modifies APP and CTF vesicular trafficking through preferential endolysosomal processing rather than autophagy.

**Figure 9:**
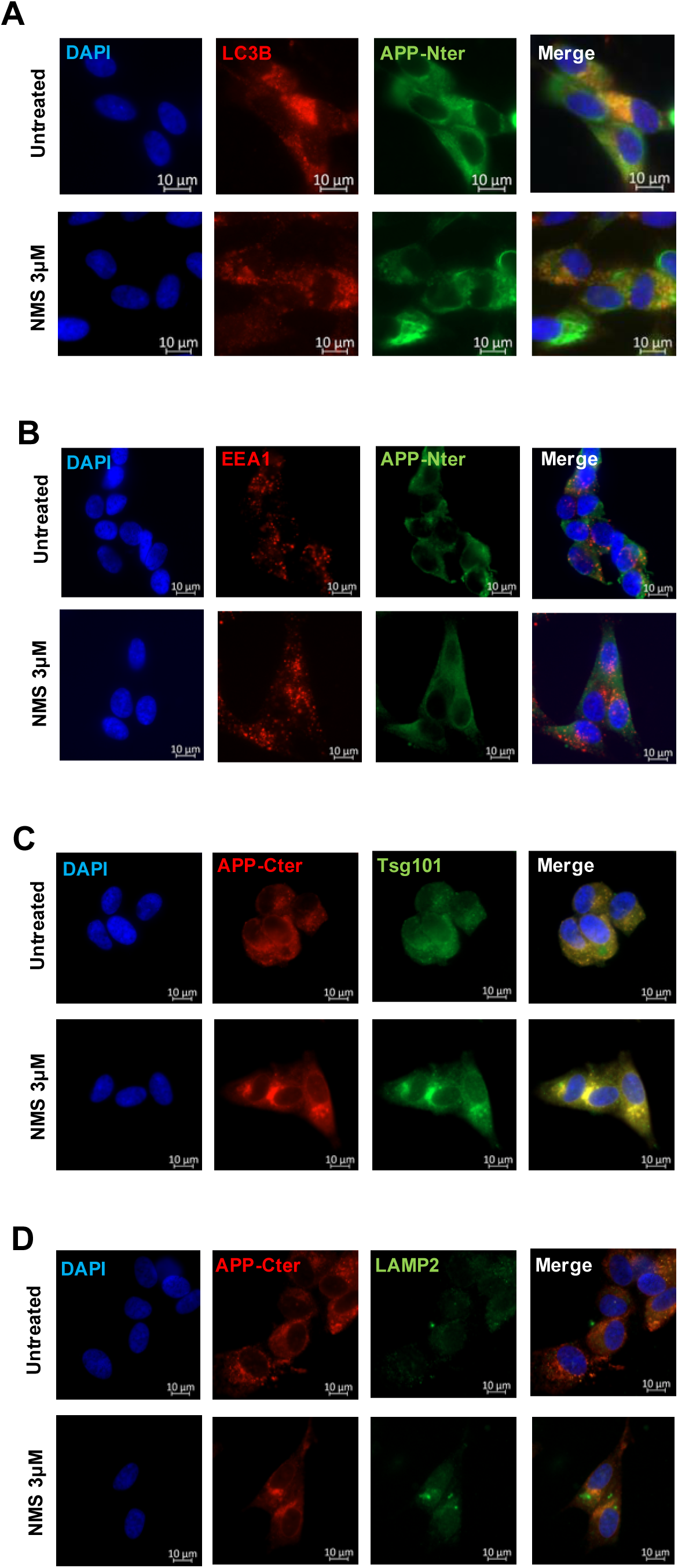
Effect of VCP ATPase inhibition by NMS-873 on APP subcellular localization in SY5Y-APP695 cells. The impact of VCP ATPase activity inhibition on APP trafficking and vesicular compartmentalization was analysed by immunofluorescence in SY5Y-APP695 cells. Cells were left untreated or treated for 24 h with 3 µM of the allosteric VCP ATPase inhibitor NMS-873, as indicated. (A) Co-immunolabeling of LC3B and APP-Nter to assess APP localization within autophagic compartments. (B) Co-immunolabeling of EEA1 and APP-Nter to evaluate APP distribution in early endosomes. (C) Co-immunolabeling of APP-Cter and TSG101 to analyze APP association with the ESCRT machinery and multivesicular body pathway (D) Co-immunolabeling of APP-Cter and LAMP2 to investigate APP localization within lysosomal compartments. Representative confocal images are shown for each condition.

## 3. Discussion

VCP is part of the AAA-ATPase family, which has various cellular functions, including breaking down misfolded proteins, protein aggregates, and damaged organelles through multiple cell-degradation pathways such as ERAD, UPS, autophagy, or endolysosomes. Several of these protein homeostasis systems influence APP metabolism. VCP may play a role in APP processing, as it has been shown to assist in degrading overexpressed βCTF (C99) via the ERAD system (Jang et al., 2017). The ribosome quality control (RQC) system manages the expression of full-length APP along the secretory pathway and involves VCP and the Hrd1 E3 ubiquitin ligase (Li et al., 2024). VCP has also been identified within the γ-secretase interactome (Wakabayashi et al., 2009). Moreover, the endolysosomal system and autophagy are reported to influence APP secretase processing and degradation, respectively (Vingtdeux et al., 2007; Tian et al., 2013). VCP also regulates the release of APP and its metabolites into extracellular vesicles, including those derived from late multivesicular endosomes (Lu et al., 2025).

Understanding APP metabolism is essential to prevent its deregulation and the buildup of toxic metabolites such as Aβ peptides or APP C99/βCTFs, which are involved in AD development (Lauritzen et al., 2012; Reiss et al., 2018). Importantly, studying APP trafficking and maturation is crucial because they both influence APP processing by secretases, as well as APP and metabolite degradation or secretion. In fact, deregulation of APP trafficking and sorting within cells is associated with changes in APP metabolism, including altered processing by secretases and degradation (Joshi and Wang, 2015; Lee et al., 2003; Wang et al., 2014, 2017).

VCP co-localizes with APP in a specific intracellular location in human and APPxPS1 transgenic brain tissue, as well as in SY5Y-APP695 and SY5Y-C99 inducible cells.

Moreover, APP and C99/βCTF are co-immunoisolated with VCP, suggesting that APP and C99 are partly part of a common protein complex. However, colocalization or co-immunoprecipitation indicates an association of VCP and APP within the same protein complex but not necessarily a direct interaction. The VCP interactome also supports the absence of a direct association, as revealed by VCP immunoprecipitation following reversible protein crosslinking in human neuroblastoma SY5Y-APP695 cells treated or not with the VCP ATPase NMS-873. APP is retrieved from the VCP interactome, along with several known APP-binding partners, including PICALM, FERMT2, SELENOS, IDE, KLC1, and NCSTN. The APP found in the VCP interactome does not distinguish between full-length protein or APP metabolites. APP-CTFs may also be co-isolated, as seen, for example, in ERAD clearance of C99, which involves SELENOS and the binding of C99 to VCP (Jang et al., 2017). From the γ-secretase complex, Nicastrin (NCSTN) was recovered, confirming that VCP interacts with one of the γ-secretase protein complexes (Wakabayashi et al., 2009), although VCP inhibition did not affect γ-secretase cleavage under our experimental conditions.

Interestingly, AD risk factors PICALM and FERMT2, both known APP interactors, were identified (Supplementary Table 1), with normalized spectra indicating increased recovery of both PICALM and FERMT2 after NMS-873 treatment.

FERMT2 interacts with APP, regulating axonal growth and synaptic plasticity through an APP-dependent mechanism (Eysert et al., 2021), whereas the PICALM/AP2 complex is suggested to act as a cargo receptor, routing endocytic APP-CTFs for autophagy degradation (Tian et al., 2013). Further studies are needed to explore the potential regulatory role of VCP in these processes.

VCP silencing and VCP ATPase inhibition were associated with changes in APP expression and processing in human neuroblastoma cells. Reducing VCP or inhibiting ATPase caused a similar accumulation of both immature and mature APP isoforms. Throughout the secretory pathway, full-length APP expression can be regulated by the ERAD or RQC system, with a key distinction: RQC pertains to protein translation, while ERAD occurs after translation. The UPS system, also known as the ubiquitin-proteasome system, primarily degrades membrane-associated unfolded or misfolded proteins. The increase in full-length APP expression was not observed with the 20S proteasome inhibitor lactacystin, indicating that proteasome inhibition does not mimic VCP inhibition. Accumulation of APP was not linked to increased APP levels during protein translation repression by cycloheximide.

Additionally, APP did not accumulate at the cell surface, indicating that VCP helps clear APP through the RQC or ERAD systems (Lu et al., 2025). When VCP is inhibited, APP bypasses these proteostasis pathways, leading to slower movement or buildup along the secretory pathway. Therefore, VCP activity may assist in sorting misfolded or unfolded APP via ERAD systems, a process that the downstream UPS does not replicate. Although speculative, these findings suggest that VCP might indirectly influence the secretory flow of APP and the levels of full-length APP along the pathway, and further research is needed to clarify this mechanism.

VCP is reported to label and prepare substrates for degradation. Therefore, loss of VCP function is thought to protect VCP substrates from being degraded. Silencing VCP or inhibiting its ATPase activity leads to the accumulation of many ubiquitinated substrates. Accordingly, full-length APP is likely a VCP substrate. In sharp contrast, under VCP silencing or ATPase inhibition, instead of an increase in APP-CTFs, APP-derived and CTF are decreased, along with C99 and C83 in SY5Y-C99 inducible cells. The use of this cell model rules out the inhibition of β-secretase, which would otherwise maintain C99 levels. Conversely, increased α- or γ-secretase activity could explain the reduced expression of APP-CTFs. VCP inhibition did not alter γ-secretase cleavage of ΔENotch, ruling out direct regulation of γ-secretase activity by VCP. Overall, our results suggest that APP-CTFs are degraded when VCP is silenced or inhibited. In fact, VCP inhibition even promotes the degradation of APP-CTFs.

VCP substrates are targeted to various proteostasis catabolic systems, including the UPS, autophagy, and the endolysosomal system. Within the interactome analyzed here, several proteins related to the early endosome, endosomal sorting machinery, and lysosome were identified, including Rab5A and Rab5B, Rab11a for recycling endosomes, and retromers VPS35 and VPS26a. Additionally, VCP’s well-established role in modulating autophagy via Beclin-1 stabilization is noted; VCP also contributes to mechanisms of the endolysosomal system, such as endosomal protein quality control and lysophagy. VCP ATPase activation by SMER28 stabilizes Beclin-1 and promotes autophagic flux. The autophagy activator SMER28 caused a reduction in APP-CTFs, similar to the effects seen with upstream mTORC1 inhibition by Torin1 or VCP inhibition. In contrast, VCP inhibition increased p62/SQSTM1 levels and did not change the LC3BII/BI ratio. Moreover, p62 silencing did not affect the profile or levels of APP-CTFs or C99-derived CTFs (supplementary Figure 6), suggesting that autophagy does not contribute to the reduction or degradation of APP-CTFs associated with VCP inhibition.

In contrast, our results suggest that the reduced expression of APP-CTFs following VCP loss of activity is mediated by the endolysosomal pathway. A mechanism that could explain the increased concentration of Aβ in SY5Y-APP695 cells through recycling endosomes (Udayar et al., 2013; Singh et al., 2025). Both BafA1 and CQ suppressed the decline of APP-CTFs caused by VCP inhibition. Through their lysosomotropic activity, they inhibit both autophagy flux and lysosome fusion with late endosomes, preventing the degradation of APP-CTFs and the production or secretion of Aβ in both scenarios. Furthermore, in SY5Y-C99 cells, although CQ and BafA1 both resulted in a reduction of C99 and C83 CTF degradation, co-treatment with a VCP inhibitor only partially prevented the degradation of C99 and C83, further indicating the role of recycling endosomal machinery. However, additional research is needed to determine the localization of APP-CTFs in endosomal compartments and autophagic vesicles.

In summary, this study identifies VCP as a novel regulator of APP metabolism. Through interactome analysis, we demonstrate that VCP and APP share a functional network that influences APP processing and trafficking in neuroblastoma cells.

Specifically, reducing VCP levels impairs APP processing, cell-surface localization, and degradation, highlighting VCP’s critical role in APP dynamics. While the precise mechanisms underlying the VCP-APP interaction remain to be elucidated, our findings establish VCP as a key modulator of APP metabolism. These results not only advance our understanding of APP regulation but also suggest VCP as a potential therapeutic target for APP-related pathologies.

### Limitation

While this study provides valuable insights into the role of VCP in APP metabolism, it is important to note that all experiments were conducted in SH-SY5Y neuroblastoma cells. Although this cell line is widely used as a model for neuronal studies, it may not fully recapitulate the physiological complexity of primary neuronal cultures or iPSC-derived neurons. Future investigations in differentiated neuronal models—such as human iPSC-derived neurons or primary cultures—will be essential to validate these findings and explore potential cell-type-specific differences in VCP-APP interactions.

## Acknowledgements

This work was supported by Inserm, Lille University, Agence Nationale de la Recherche (ANR): LabEx DISTALZ ANR-11-LABX-01, ANR VIDALZ ANR-15-CE18-0002, France Alzheimer. Adriana Figueroa-Garcia is a recipient of a postdoctoral fellowship from the Alzheimer’s Disease Association, France. Catherine Baud was a postdoctoral fellow of the ANR VIDALZ. Alexandre Trichies was an engineer for the ANR VIDALZ. Caroline Evrard was a recipient of a doctoral fellowship from Lille University. Lisa Brisoire is a recipient of a fellowship from Lille University.

## Author contributions

AFG, LB, CB, RC, SE, and CE contributed to all cell experiments. MC and SB developed the SY5Y-APP695 clones. CB and JMS conducted the VCP interactome analysis. PM headed the ANR VIDALZ, and Luc BUEE headed the France Alzheimer VCP project. CE and VV performed the analysis of the secretory pathway. AFG and LB analyzed APP and C99 metabolism. RPP and FC contributed to manuscript writing. NS designed the entire project, analyzed the results, and authored the manuscript.

## Ethics approval and consent to participate

All applicable international, national, and institutional guidelines for the care and use of animals were followed. Experimental procedures involving animals were approved by the Lille Neuroscience and Cognition Research Center, Alzheimer & Tauopathies, and Lille Neuro & Chemistry teams, as well as the Use Committee and the National Council on Animal Care of the Ministry of Health (Project No. APAFIS# 2023050714441306 and APAFIS #49656-2024020215104250), in accordance with the guidelines established by the European Community Council (Directive of November 24, 1986) and the ethical standards of the institutional animal welfare committee.

All procedures involving human participants were conducted in accordance with the ethical standards of the institutional and national research committees and with the principles of the 1964 Declaration of Helsinki and its subsequent amendments or comparable ethical standards. Human brain samples were obtained from the Lille NeuroBrain Bank (CRB/CIC1403). Written informed consent was obtained from the patients themselves or, where applicable, from their next of kin, in accordance with French bioethical regulations (authorization AC-2013-1887). All cases were anonymized; however, information regarding sex, age at death, and neuropathological diagnosis was provided.

---

**Supplementary figure 1:**
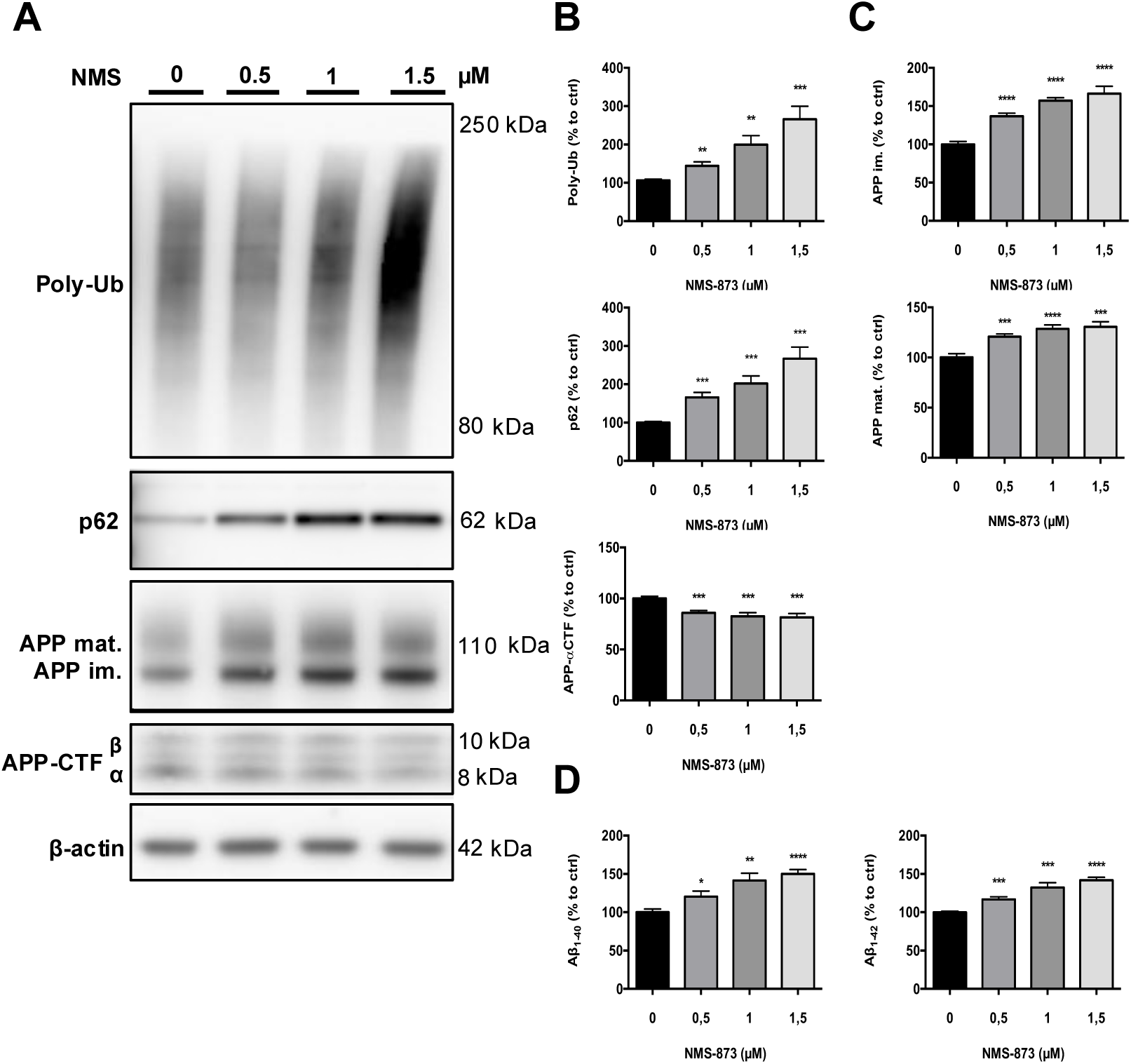
Effect of VCP ATPase activity inhibition by NMS-873 on APP metabolism in SY5Y-APP695 cells. The inhibition of VCP recombinant ATPase activity was tested using increasing concentrations of the allosteric VCP ATPase inhibitor NMS-873, and the IC_50_ was determined (A). SY5Y-APP695 cells were treated for 24 hours with increasing concentrations of NMS-873, as indicated. Cell lysates were immunolabeled with the following antibodies: Poly-ubiquitinylated proteins (Poly-Ub), APP-Cter-C17, and β-actin as a loading control (B). Densitometric analysis and quantification of VCP, Poly-Ub proteins, immature and mature APP levels, and APP ⍺-carboxy-terminal fragments were performed (C and D). ELISA analysis of the conditioned medium was used to measure the concentration of secreted Aβ_1-40_ and Aβ_1-42_ peptides (E). Histograms indicate the mean ± SD. n=6 *p<0.05, **p<0.01, ***p<0.005, ****p<0.001 Unpaired *t*-test.

**Supplementary figure 2:**
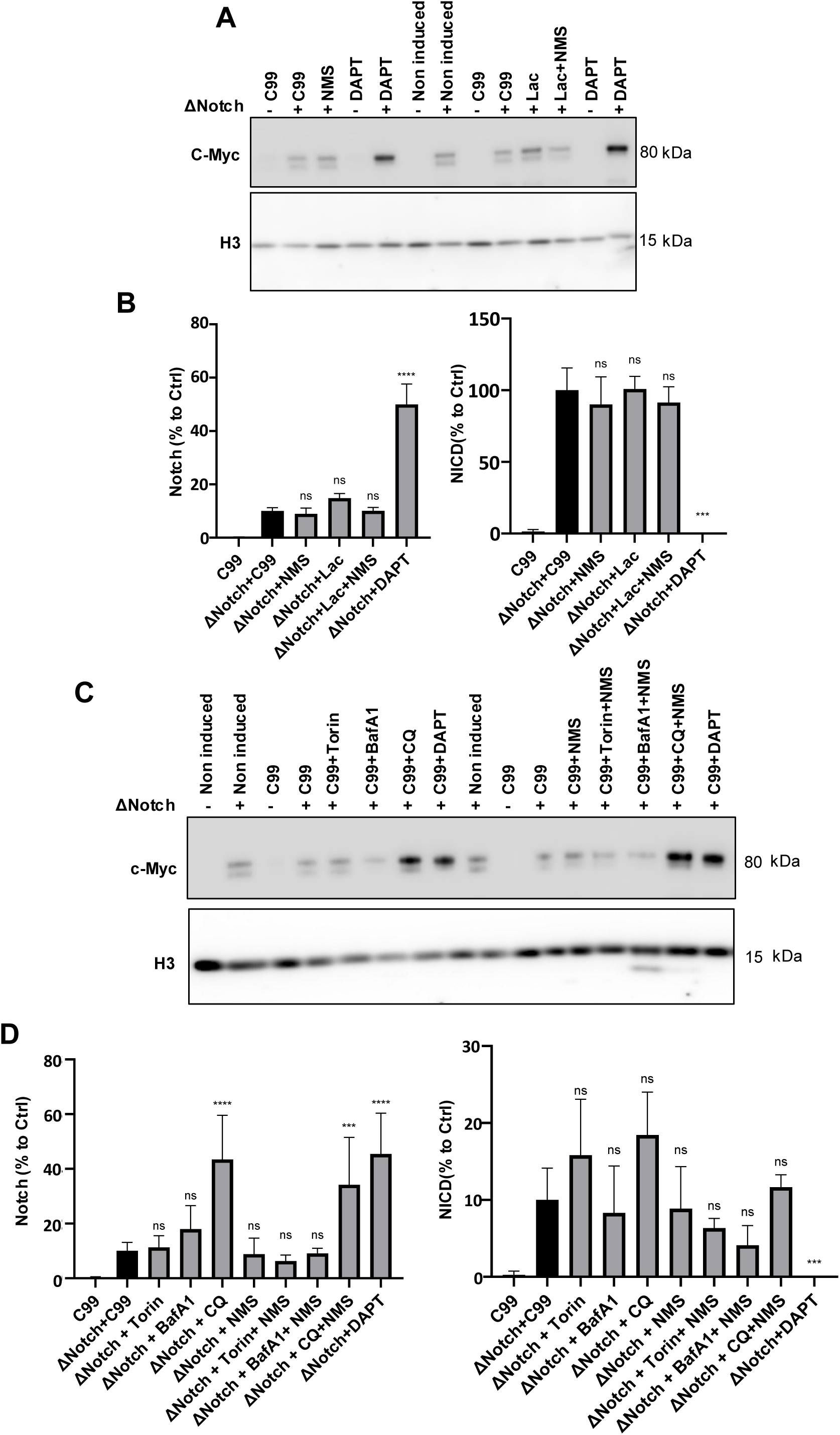
Gamma-secretase cleavage of ΔENotch is unchanged by autophagy activation or inhibition and NMS-873 treatments. (A-C) Immunoblot analysis of Notch-cleaved products by γ-secretase in SY-C99 cells. Extracts of SY-C99 cells transfected with NotchΔE were detected with the anti-myc antibody. Histone H3 was used as a loading control. Some lanes were treated with 1,5 µM of NMS-873, 1µM of Lactacystin, Torin1 1 µM, 30 µM Chloroquine,100 nM Bafilomycin A1 and 200 nM DAPT. (B-D) Quantification, data are expressed as the mean ± SD, (n = 3 independent experiments), ns=non-significant.

**Supplementary figure 3:**
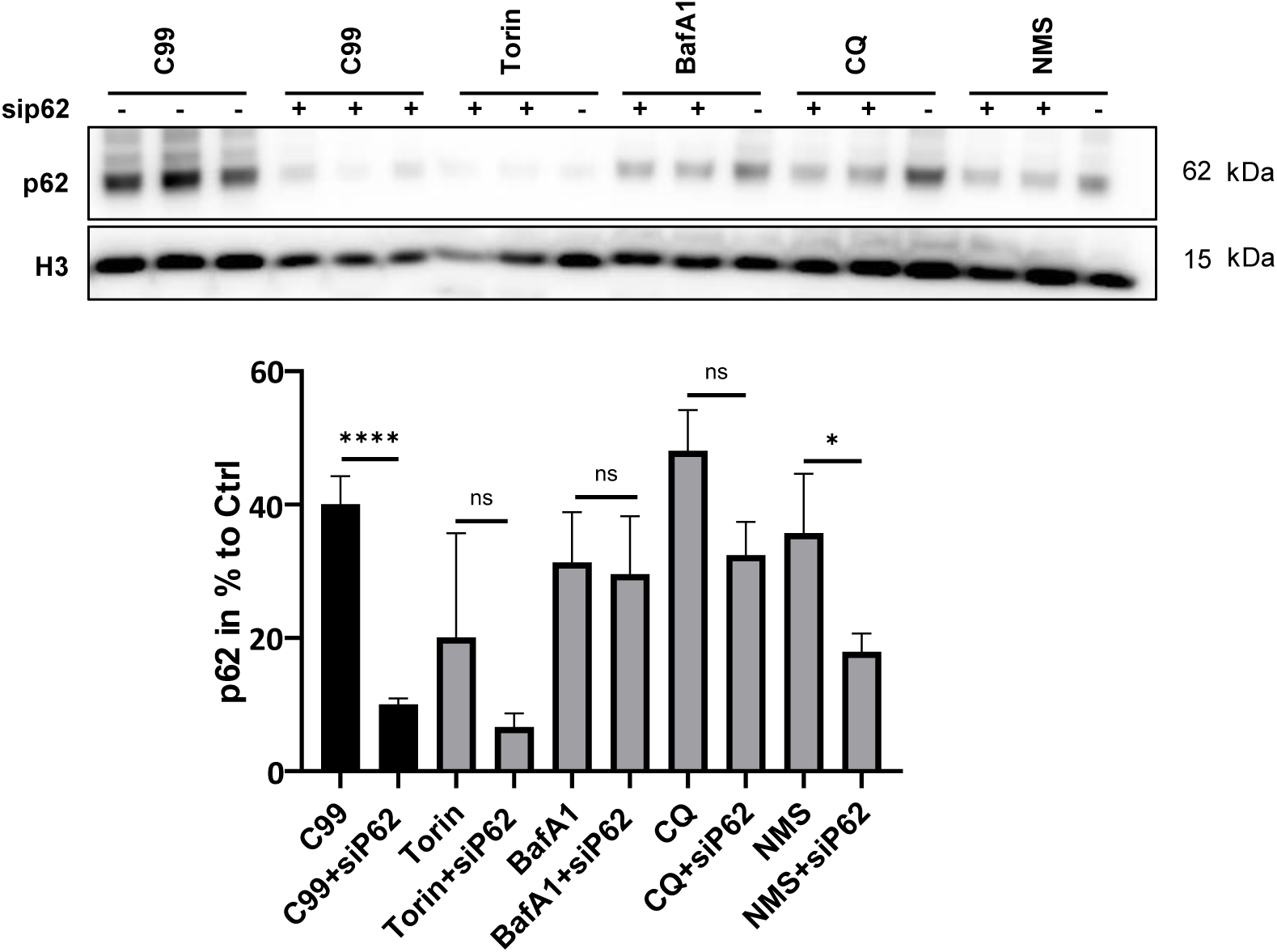
**Validation of p62 siRNA in SY5Y-APP695 and SY5Y-C99 cells**. Cells were transfected with p62 siRNA (sip62) for 72 hours. Cell lysates were immunolabeled with antibodies targeting p62 and H3 as a loading control (A). Densitometric analysis and quantification of p62. Histograms represent the mean ± SD. n=4; ns=non-significant, *p<0.05, ****p<0.001 Unpaired t-test.

**Supplementary figure 4:**
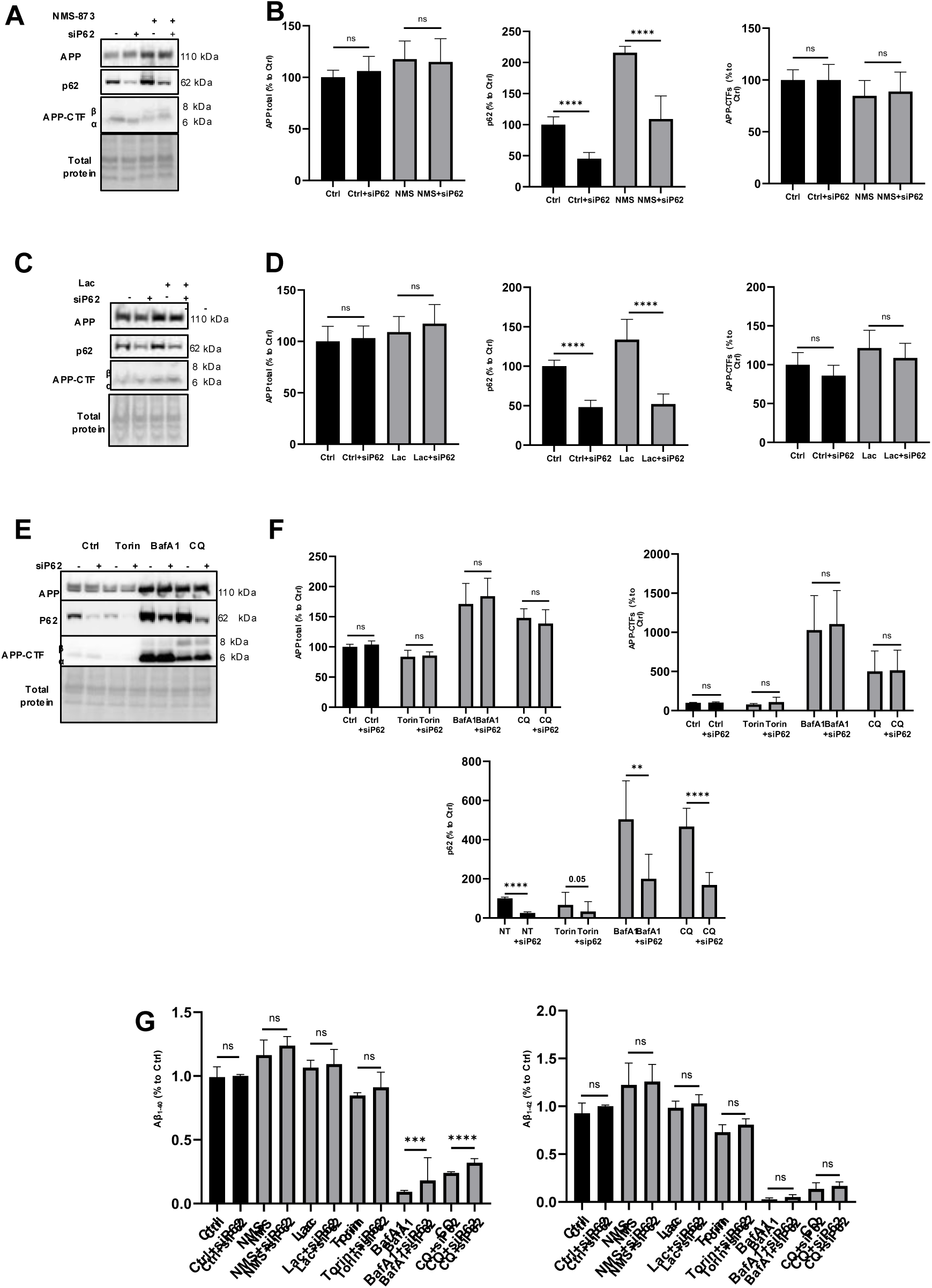
Effect of p62 siRNA on APP metabolism in SY5Y-APP695 cells. SY5Y-APP695 cells were transfected with p62 siRNA (siP62) for 72 hours. Cell lysates were immunolabeled with antibodies targeting p62, APP-Cter-C17, and Total protein load as a loading control (A, C, E). Densitometric analysis and quantification of p62, APP levels, and APP α-carboxy-terminal fragments (B, D, F). The concentrations of Aβ1-40 and Aβ1-42 peptides were measured in the cell conditioned media by ELISA (G). Histograms represent the mean ± SD. n=3 ns=non-significant, **p<0.01, ***p<0.005, ****p<0.001 Unpaired t-test.

